# Wax ester synthase overexpression affects stomatal development, water consumption and growth of poplars

**DOI:** 10.1101/2024.04.06.588381

**Authors:** Ashkan Amirkhosravi, Gerrit-Jan Strijkstra, Alisa Keyl, Felix Häffner, Ulrike Lipka, Cornelia Herrfurth, Ivo Feussner, Andrea Polle

**Affiliations:** Forest Botany and Tree Physiology, Büsgen-Institute and Göttingen Center for Molecular Biosciences (GZMB), Büsgenweg 2, University of Göttingen, 37077 Göttingen, Germany; Department of Plant Biochemistry, Albrecht-von-Haller-Institute of Plant Sciences and Göttingen Center for Molecular Biosciences (GZMB), Justus-von-Liebig-Weg 11, Universitäty of Göttingen, 37077 Göttingen, Germany; Service Unit for Metabolomics and Lipidomics, Göttingen Center for Molecular Biosciences (GZMB), Justus-von-Liebig-Weg 11, University of Göttingen, 37077 Göttingen, Germany

**Keywords:** Drought stress, wax ester synthase, stomatal conductance, water use efficiency, cuticle wax, occluded stomata

## Abstract

Poplars are important fast-growing biomass crops. Their water-spending lifestyle renders them susceptible to drought and threatens plantations under global climate change with extended periods of water deprivation. The cuticle and stomatal regulation are major traits to protect plants from uncontrolled water loss. Here, we targeted the wax biosynthesis pathway of *Populus* x *canescens* by overexpressing jojoba (*Simmondsia chinensis*) wax ester synthase (Sc*WS*) to improve cuticular properties. Sc*WS* expression caused accumulation of lipid droplets inside the cells, decreased transcript levels of endogenous wax biosynthetic genes, and moderate shifts in surface wax composition but did not affect non-stomatal water loss. During short- and long-term drought scenarios under greenhouse and outdoor conditions, Sc*WS* lines showed decreased stomatal conductance and increased water-use-efficiencies leading to a water-saving phenotype and delayed leaf shedding. This phenotype was caused by a high fraction (80%) of wax-occluded or semi-occluded stomata, and was accompanied by suppression of *OCCLUDED STOMATAL PORE1* (*OSP1*), known to cause abberant wax accumulation at the stomatal ledges as found here. Occluded stomata limited poplar photosynthesis under high but not under low light intensities. Leaf damage and insect scores did not reveal differences compared with wild-type plants. Biomass production of Sc*WS* lines was unaffected in short-term experiments but dropped below that of wild-type poplars at the end of two field seasons, indicating a growth trade-off. In conclusion, our study pinpoints a tight connection between wax biosynthesis and stomatal features and opens a new avenue to improve poplar water consumption by optimizing stomatal ledges with refined biotechnological approaches.

## Introduction

Climate change is predicted to result in more frequent and intense droughts as a consequence of anthropogenic impacts (Trenberth *et al*., 2014; Chiang *et al*., 2021). Excess tree mortality is linked to drought events (Senf *et al*., 2020) and can be a direct result of hydraulic failure or occurs in the course of secondary incidents such as pathogen and insect calamities that kill stressed trees (Anderegg *et al*., 2013; Jactel *et al*., 2012). It is therefore important to explore the physiological mechanisms that can be exploited to better protect trees against water loss during drought events.

The evolutionary adaptation of plants to live in terrestrial habitats required strict control of the water status (Xue *et al*., 2017). Water flux through the plant system and uptake of atmospheric CO_2_ for photosynthetic assimilation are regulated by stomata; their frequency, size, and responsiveness to changes in water availability affect plant drought performance (Jeffree *et al*., 1971). Further, plant surfaces are covered by a hydrophobic layer, the cuticle, which drastically reduces surface evaporation and constitutes a barrier against biotic invaders (Lewandowska *et al*., 2020). The cuticle contains, among other compounds, cutin, and waxes, whose composition varies with environmental conditions (Lewandowska *et al*., 2020; Shaheenuzzamn *et al*., 2021). Together, stomatal features and cuticular properties are major traits that determine plant drought resistance.

The composition and production of the cuticle as a shield against water loss has received increasing attention (Samuels *et al*., 2008; Zeisler-Diehl *et al*., 2020). Waxes are water-repellent and mainly consist of very long-chain (VLC) aliphatics and specialized metabolites, such as terpenoids and phenylpropanoids (Busta and Jetter, 2018; Jetter and Riederer, 2016). Differential removal of these classes showed that VLC aliphatics are the principal constituents blocking cuticular water loss (Seufert *et al*., 2022).

The wax biosynthesis starts with the elongation of C16 and C18 acyl-Coenzyme As (CoAs) by ER-associated, multimeric FATTY ACID ELONGASE (FAE) complexes, which sequentially extend the acyl-CoA chains by two carbon atoms per elongation cycle (Lewandowska *et al*., 2020). For wax production, the acyl-CoAs are elongated to VLCFAs between 20 and 38 carbons in length (Samuels *et al*., 2008; Hegebarth *et al*., 2017). VLC acyl-CoAs can be either exported as free VLCFAs or undergo further modifications via the alcohol- or alkane-forming pathway (Lewandowska *et al*., 2020). In the alcohol-forming pathway of *Arabidopsis thaliana*, VLC acyl-CoAs are reduced to primary alcohols by the fatty acyl-CoA reductase 3 (*At*FAR3/ECERIFERUM 4 [*At*CER4]). For the production of wax esters, primary alcohols are esterified with mainly C16 acyl-CoAs by the bifunctional wax ester synthase/acyl-CoA:diacylglycerol acyltransferase 1 (*At*WSD1s) (Li *et al*., 2008). Further analyses about the role of *At*WSD1 in abiotic stress responses uncovered that *At*WSD1 promotes drought tolerance (Patwari *et al*., 2019). Moreover, heterologous *AtWSD1* expression in *Camelina sativa* caused a massive increase in the thickness of the cuticle wax layer and mediated enhanced resistance against various osmotic stresses (Abdullah *et al*., 2021). In addition to bifunctional WSDs, mono-functional acyltransferases (WS) are also involved in plant wax biosynthesis. Among these enzymes, wax ester synthase from the drought-tolerant desert species jojoba *Simmondsia chinensis* (*Sc*WS) is of particular interest (Lardizabal *et al*., 2000). The jojoba waxes are composed of monoenoic aliphatic chains of high stability; they are used for cosmetic products and in addition have an enormous potential for industrial applications since they can be produced by bioengineering as feedstock for lubricants, biodiesel, etc. (Iven *et al*., 2016; Yu *et al*., 2018; Alotaibi *et al*., 2020; Gad *et al*., 2021). Whether *Sc*WS can also contribute to enhancing drought resistance of biomass/bioenergy plants and thus, provide dual benefits, is unknown.

Here, we used poplar (*Populus* sp.) trees to clarify the role of *Sc*WS in mediating drought resistance. Poplars are economically important biomass crops because of their fast growth and wood production (Taylor *et al*., 2019). However, their productivity is often limited by water availability, and therefore, massive attempts are underway to improve poplar drought resistance (Polle *et al*., 2019). In recent studies, transcription factors regulating cuticular wax biosynthesis were targeted to induce drought tolerance in poplars. For example, overexpression of *AtWIN1/SHN1* in *Populus* x *canescens* resulted in glossy leaves, decreased stomatal densities, and improved water-use efficiency (Lawson, 2017). Similarly, *PeSHN1* from *Populus × euramericana* overexpressed in *Populus alba × P. glandulosa* increased wax accumulation and improved drought performance (Meng *et al*., 2019). Members of the MYB transcription factor family (MYB94, MYB142, MYB96) also play a role in the regulation of cuticular wax biosynthesis and drought responses (Seo *et al*., 2011; Cui *et al*., 2016; Yin *et al*., 2017). The deregulation of the transcription factors causes changes in the composition of the cuticular waxes, and cuticular thickness and affects stomatal conductance (Fang *et al*., 2020; Song *et al*., 2022). However, in many plant species, including poplar, the significance of the thickness of the cuticule for drought tolerance has been questioned because its variations did not affect the water permeability of leaves (Grünhofer *et al*., 2021; Grünhofer *et al*., 2022), while the wax composition may be crucial for water retention. For example, transgenic suppression of β-ketoacyl-CoA synthase *CER6*, a subunit of the acyl-CoA elongase complex in poplar resulted in an increased chain of wax esters, which did not affect the total wax load but increased non-stomatal water loss (Grünhofer *et al*., 2024). Considering the mixed and partly overlapping or conflicting effects of biosynthetic and regulatory genes on plant water balance, a more comprehensive analysis of wax biosynthesis on stomatal and cuticular traits is needed to improve poplar fitness under global change conditions.

Here, we expressed *Sc*WS in *Populus* x *canescens.* We expected that the mono-functional jojoba wax ester synthase would increase the hydrophobicity of the cuticle, thereby, reducing the evaporation of water and improving the fitness of the transgenic plants under drought. We tested the performance of wild type (WT) and *Sc*WS poplars under greenhouse conditions and in small plantation-like settings under outdoor conditions for two growth seasons to explore their biotechnological potential. We show that expression of *Sc*WS increases the cellular lipid content, and has moderate effects on wax composition but does not augment the overall cuticular wax load of the leaf. Under these conditions, the stomatal development, especially the formation of stomatal ledges was affected by an aberrant wax accumulation on their surface, resulting in an occluded stomatal phenotype. Consequently, *Sc*WS poplars had lower stomatal conductance and lower photosynthesis but higher water-use efficiency under greenhouse and long-term outdoor conditions. We also investigated the possible ecological consequences of *Sc*WS overexpression for insect interactions and wood biomass production. Together our data show a tight connection between wax biosynthesis and stomatal features.

## Results

### Characterization of *Sc*WS-expressing poplar lines reveals a stomatal phenotype

Poplar (*Populus* x *canescens*) transformed with the jojoba *Sc*WS under the *35S* promoter (called *Sc*WS lines) showed high expression levels of the transgene (Fig 1a). Since *AtWSD1* is responsible for wax ester production of the cuticle of *A. thaliana* (Li *et al*., 2008), we conducted a phylogenetic analyses of poplars *WSD*s (Supplement Figure S1, conducted according to Cheng *et al*., 2022) to identify close homologues of *AtWSD1*. We chose *Pc*WSD1 as the reference in WT plants because its expression level was not affected by Sc*WS* expression (Fig. 1b), whereas *Pc*WSD4, the closest homologue to *At*WSD1, showed drastically suppressed transcript level in the *Sc*WS lines compared with the WT (Fig. 1c).

**Figure 1.**
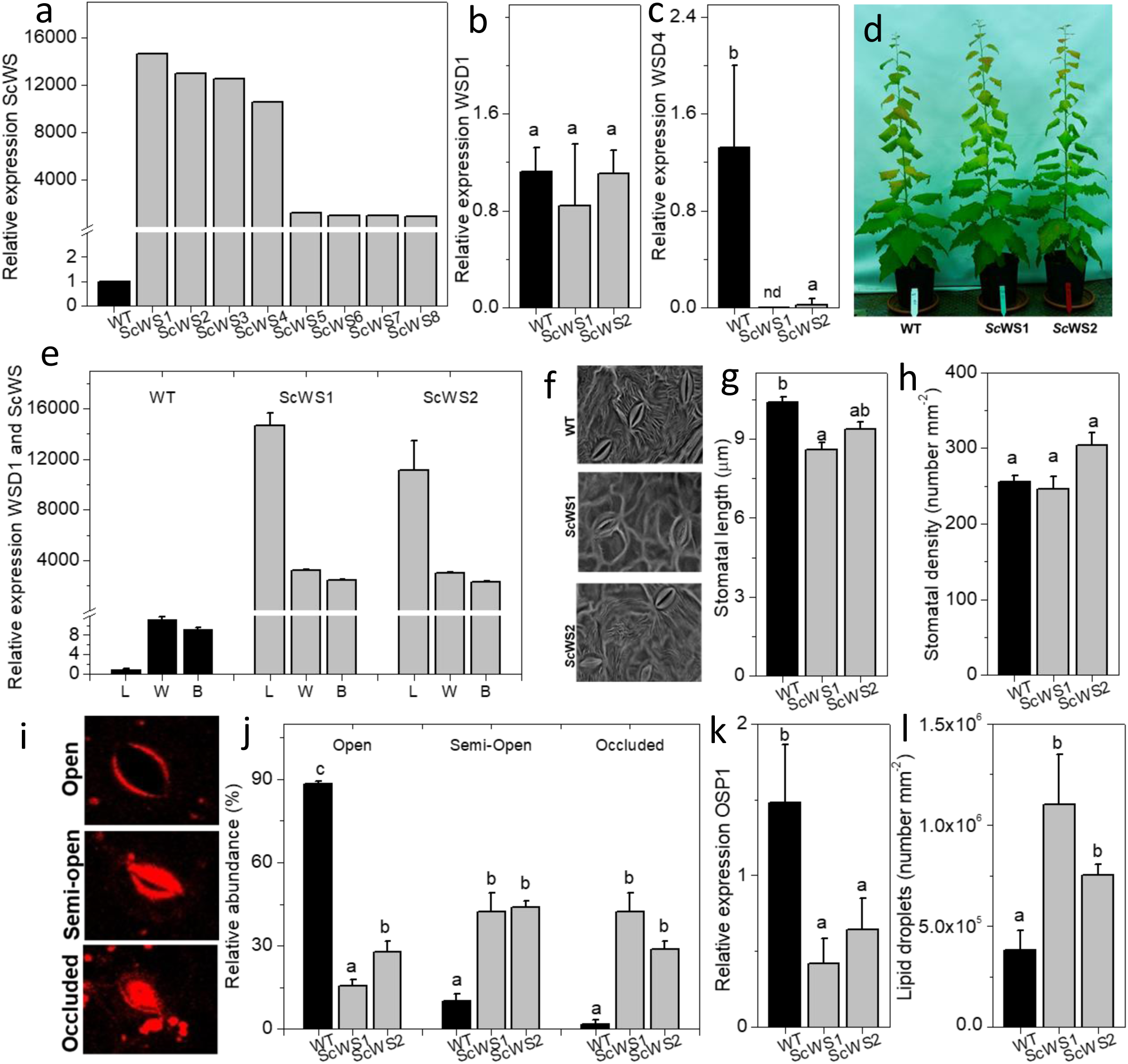
Characterization of wax ester synthase (*ScWS*) expressing lines compared with wild-type *P*. x *canescens* (*Pc*). Black bars indicate wild type (WT) and grey bars *ScWS*-expressing lines. (a) Expression of *ScWS* under the *35S* promoter in several independent poplar lines, (b) expression of *PcWSD1*, (c) *PcWSD4*, and (d) phenotype of WT and *ScWS* expressing lines, (e) expression of *PcWSD1* and *ScWS* in leaf, wood and bark. The expression of *ScWS* was normalized to *WSD1*, nd = not detected. (f) scanning electron microscopy of the lower leaf surface showing cuticular ridges emerging from the stomata and lack of ridges in the lines *ScWS1* and *ScWS2*, (g) stomatal lengths, (h) stomatal density, (i) fluorescence microscope of typical stomatal anatomy of the *ScWS* lines; images were recorded after staining with the lipophilic dye Lipid spot II, (j) quantification of stomatal types. The total number of counted stomatal was set as 100%, (k) expression of *PcOSP1*, (l) number of lipid droplets per mm² in WT and *ScWS* lines. Lipid droplets were counted in four areas of 10 x 10 µm² per indididual sample and normalized to 1 mm². For the selection of genotypes with high *ScWS* expression one leaf per line was analysed. Further analyses were conducted with 4 to 7 individual samples per line. Data show means (± SE). Different letters indicate significant differences among means at *p* ≤ 0.05 (ANOVA, *post-hoc*Tukey test).

The growth phenotype of *Sc*WS lines was similar to that of WT plants (Fig. 1d). Sc*WS* was expressed in all inspected poplar tissues (leaf, bark, wood, Fig. 1e). The suppression of Pc*WDS4* in the transgenic *Sc*WS lines was specific to leaves and bark, and not found in poplar wood or developing xylem (Supplement Fig. S2).

Leaf cross sections inspected by transmission electron microscopy (TEM) did not reveal changes in the thickness of the cuticular layer (Supplement Fig. S3) but scanning electron microscopy showed changes in the patterns of crystal-like wax structures on the upper leaf surface (Supplement Fig. S4) and partial disappearance of cuticular ridges on the lower leaf surface (Fig. 1f, Supplement Fig. S5). Furthermore, the stomatal lengths of the *Sc*WS lines were shorter than those of the WT (Fig. 1g) but the stomatal frequencies were similar (Fig. 1h).

Since the stomatal features can be affected by environmental conditions, we asked if the stomatal development of the transgenic lines was more susceptible than that of the WT to fluctuations in temperature or humidity under greenhouse cultivation. However, stress indicators such as abscisic acid were unaffected in the *Sc*WS lines compared to the WT cultivated under greenhouse conditions (Supplement Fig. S6). Transcript levels of key transcription factors involved in the regulation of wax biosynthesis in stressed plants (Seo *et al*., 2011) were also unaffected (*PcMYB94*) or even decreased (*PcMYB96*) in the *Sc*WS lines compared with the WT (Supplement Fig. S6). This result speaks against increased susceptibility to ambient environmental conditions in the *Sc*WS lines. Furthermore, we inspected *Sc*WS and WT plantlets grown under high humidity in closed sterile tissue culture jars to exclude environmental stress (Supplement Fig. S7). Under these conditions, the stomata of the *Sc*WS lines were also smaller than those of empty vector controls and the WT (Supplement Fig. S7). Therefore, we conclude that the shorter stomatal phenotype was a genetic consequence of heterologous Sc*WS* expression.

Close inspection of the stomata by TEM and confocal laser scanning microscopy (CLSM) revealed that 40% were semi-occluded and 30% to 40% were occluded by excess wax accumulation in the *Sc*WS lines compared to the WT (Fig. 1i,j). The accumulation occurred mainly at the stomatal ledge (Fig. 1i). The abberant wax deposition caused a stomatal phenotype that strongly resembled that reported by Tang *et al*. (2020) in an *A. thaliana* null mutant of *OSP1* (*OCCLUDED STOMATAL PORE 1*). Therefore, we measured the relative transcript abundance of *PcOSP1* in WT and *Sc*WS lines. We found that the expression levels of *PcOSP1* were drastically decreased in the *Sc*WS lines compared with the WT (Fig. 1k), thus, supporting a link between the wax ester biosynthesis and stomatal development. In addition to the occluded stomatal phenotype, we noted a massive accumulation of lipid droplets in the *Sc*WS lines compared with the WT (Fig. 1l, Supplement Fig. S8). Thin layer chromatography of whole-leaf extracts also showed an accumulation of wax esters in *Sc*WS lines compared with WT plants (Supplement Fig. S9).

### *Sc*WS modulates the biosynthesis of surface waxes under drought

We studied the expression of *Sc*WS and the biosynthetic key genes involved in the alcohol-forming pathway leading to wax esters and in the alkane-forming pathway required for the production of VLC aldehydes, alkanes and alkenes under well-irrigated and drought-stressed conditions (Fig. 2). Both pathways start with the elongation of fatty acids (C16/C18 acyl CoA) including among other enzymes, CER2 and CER6 (Lewandowska *et al*., 2020). Under non-stressed conditions, the transcript levels of *CER2* and *CER6* were not significantly affected in the *Sc*WS lines compared with the WT but showed a declining trend (Fig. 2a,b). Under drought however, the transcript levels increased. The increase was generally more pronounced in the WT than in the *Sc*WS lines (Fig. 2a,b)

**Figure 2.**
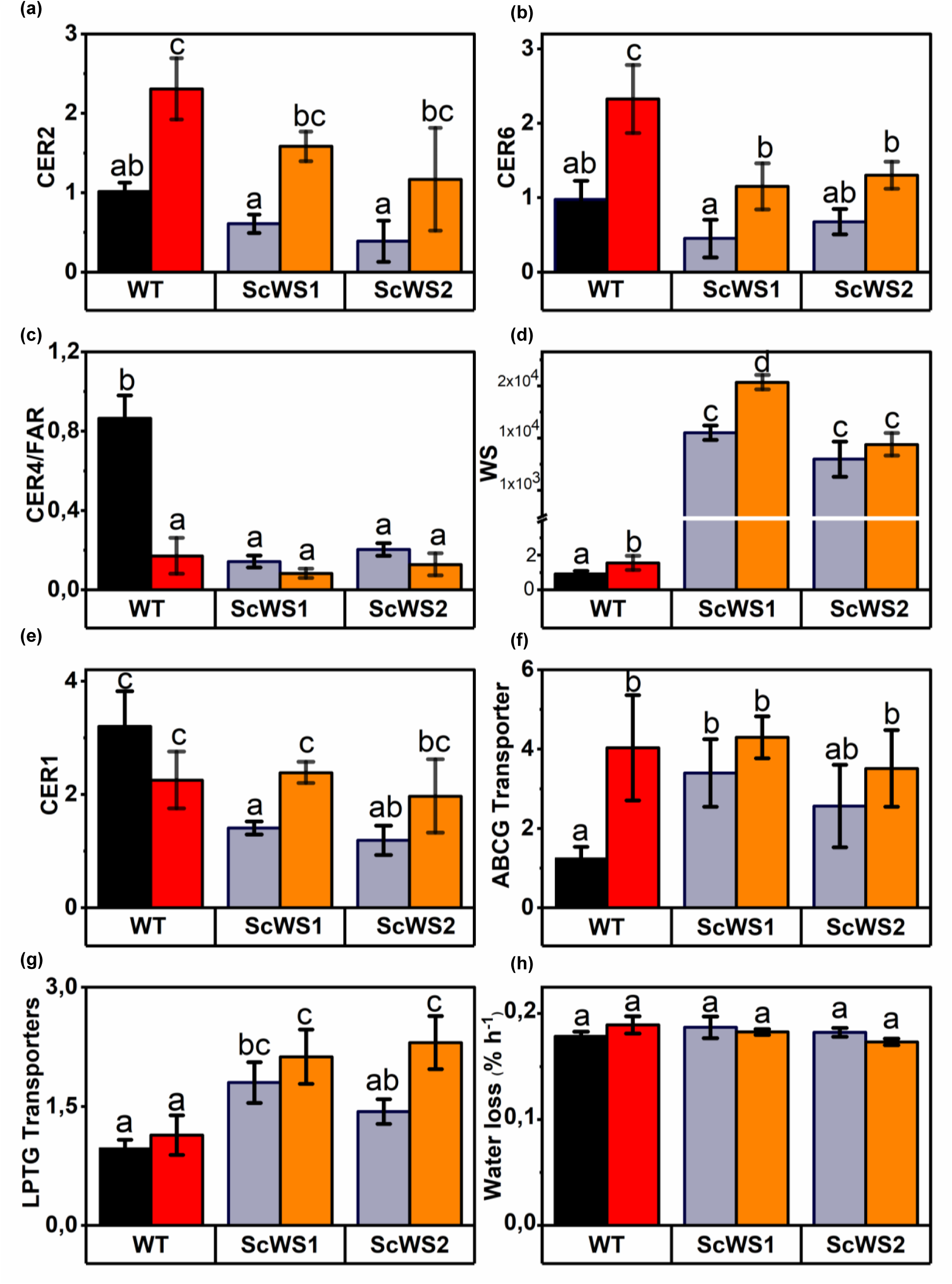
Relative expression levels of genes involved in wax biosynthesis in wild-type (WT) and *ScWS* expressing poplars under well-irrigated and drought-stressed conditions. (a) *CER2* and (b) *CER6*, which are potentially involved in the elongation of fatty acyl CoA to very long chain (VLC) fatty acyl CoA, (c) *FATTY ACID REDUCTASE* (*CER4/FAR3*) which provides the precursors for wax ester synthase, (d) wax ester synthase (ScWS) in transgenic lines and *WSD1* in the WT, (e) *CER1*, which diverts VLCFA into the alkane/alkene pathway, (f) putative lipid transporters of the adenosine triphosphate binding cassette family (ABCG) and (g) of the lipid transfer protein family (LTPG), (h) velocity of non-stomatal water loss. Black = WT well-irrigated, grey = *ScWS* lines well-irrigated, red = WT drought-stressed, orange = *ScWS* lines drought stressed. Control and stressed leaves were harvested after 21 days of withholding water from the long-term drought experiment under greenhouse condtions. Data show means (n = 4 individual plants per line and treatment ± SE). Data were log-transformed, analysed by ANOVA and the *post-hoc* Tukey test. Different letters indicate significant differences among the treatments and lines at *p* ≤ 0.05.

The reduction of VLC acyl-CoAs by CER4/FAR3 is the first committed step in the alcohol-forming pathway, followed by wax ester synthesis via WSDs. Under non-stressed conditions, the transcript levels of Pc*CER4/FAR3* were about 9-fold lower in the *Sc*WS lines than in the WT (Fig. 2c). Under drought, the Pc*CER4/FAR3* levels decreased in the WT and remained low in the *Sc*WS lines (Fig. 2c). The Pc*WSD1* transcript levels increased two-fold in WT leaves in response to drought but were still more than three orders of magnitude lower than those of the constitutively expressed Sc*WS* in the transgenic lines (Fig. 2d).

The VLC acyl-CoAs are converted into VLC aldehydes, alkanes, and alkenes in the alkane forming pathway. The production of alkanes is promoted by CER1, synergistically with other enzymes (Aarts *et al*., 1995; Bourdenx *et al*., 2011). In WT poplars, *PcCER1* transcript levels were not significantly affected by drought (Fig. 2e). In the non-stressed *Sc*WS lines, the transcript levels of *PcCER1* were about twice lower than in the WT but increased to WT levels in response to drought (Fig. 2e).

Transporters for lipophilic compounds to the leaf surface (ABCG, LTPG) (Pighin *et al*., 2004; DeBono *et al*., 2009) were increased by Sc*WS* expression as well as under drought (Fig. 2f,g). This result does not support restrictions on the export of aliphatic compounds.

Despite massive Sc*WS* expression in the transgenic lines, the water loss assay, which indicates the velocity of non-stomatal water loss, did not reveal significant differences compared with WT leaves (Fig. 2h).

We scrutinized the influence of Sc*WS* expression on cuticular wax composition and total wax load under well-irrigated and drought-stressed conditions (Fig. 3). The wax composition of the WT consisted of fatty acids, primary alcohols, wax esters, aldehydes, alkanes, and alkenes (Fig. 3a), being similar to that reported previously for *P.* x *canescens* (Grünhofer *et al*., 2021). The profile of VLC aliphatics and wax esters of the *Sc*WS lines was similar to that of the WT but contained significantly less C27 alkanes and C28 primary alcohols, which were the major constituents of the cuticle (Fig. 3b,c). Some of the minor compounds in the group of alkanes, primary alcohols, and wax esters were also slightly diminished compared with WT (Fig. 3). The differences in wax load among the WT and the ScWS lines disappeared under drought (Fig. 3d,e,f), mainly because of elevated levels of C27 alkanes and C28 primary alcohols in the *Sc*WS lines) (Fig. 3b,c).

**Figure 3.**
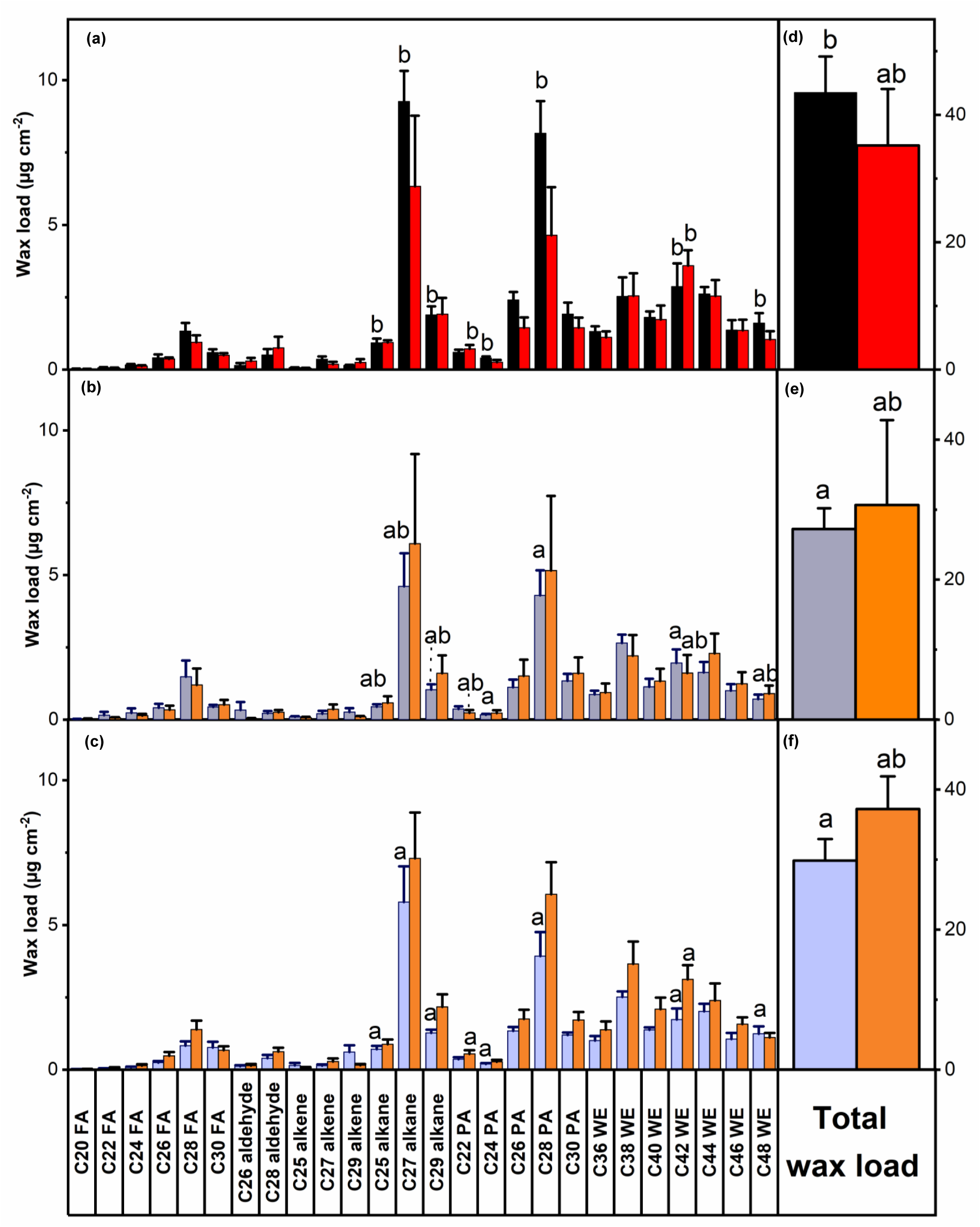
Composition and amounts of cuticle waxes of *P*. x *canescens* leaves of the wild type (WT) (a,d) and transgenic lines *ScWS1* (b,e) and *ScWS2* (c,f). Cn = carbon chain chain lengths, FA = fatty acid, PA = primary alcohol, WE = wax esters. Panels a,b,c show the abundance of the individual compounds and panels d,e,f the total wax load as the sum of all compounds. Bars show means (± SE) with n = 5 biological replicates per line and treatment (exceptions: WT control n = 4, *Sc*WS2 drought n = 3). ANOVA, post-hoc least significant difference test. Different letters across the panels indicate significant differences at *p* ≤ 0.05 among the lines for an individual compound. Black = WT well-irrigated, grey = *ScWS* lines well-irrigated, red = WT drought-stressed, orange = *ScWS* lines drought stressed.

### Sc*WS*-expressing poplars exhibit enhanced water use efficiency but long-term growth penalty

Under well-irrigated conditions (soil moisture > 0.3 m³ m^-^³), the stomatal conductance of WT plants ranged from 600 to 800 mmol m^-2^ s^-1^ and was almost two times higher than that of the *Sc*WS lines (Fig. 4a). Under drought (soil moistures of about 0.1 m³ m^-^³), the stomatal conductance of the WT lines declined to levels similar to those of the non-stressed transgenic lines (Fig. 4a). These response patterns were found in different independent experiments by withholding water for increasing drought periods from one week (short-term greenhouse) to three weeks (long-term greenhouse) or almost two months under field conditions in a plantation-like growth array (Fig. 4a, Supplement Fig. S10).

**Figure 4.**
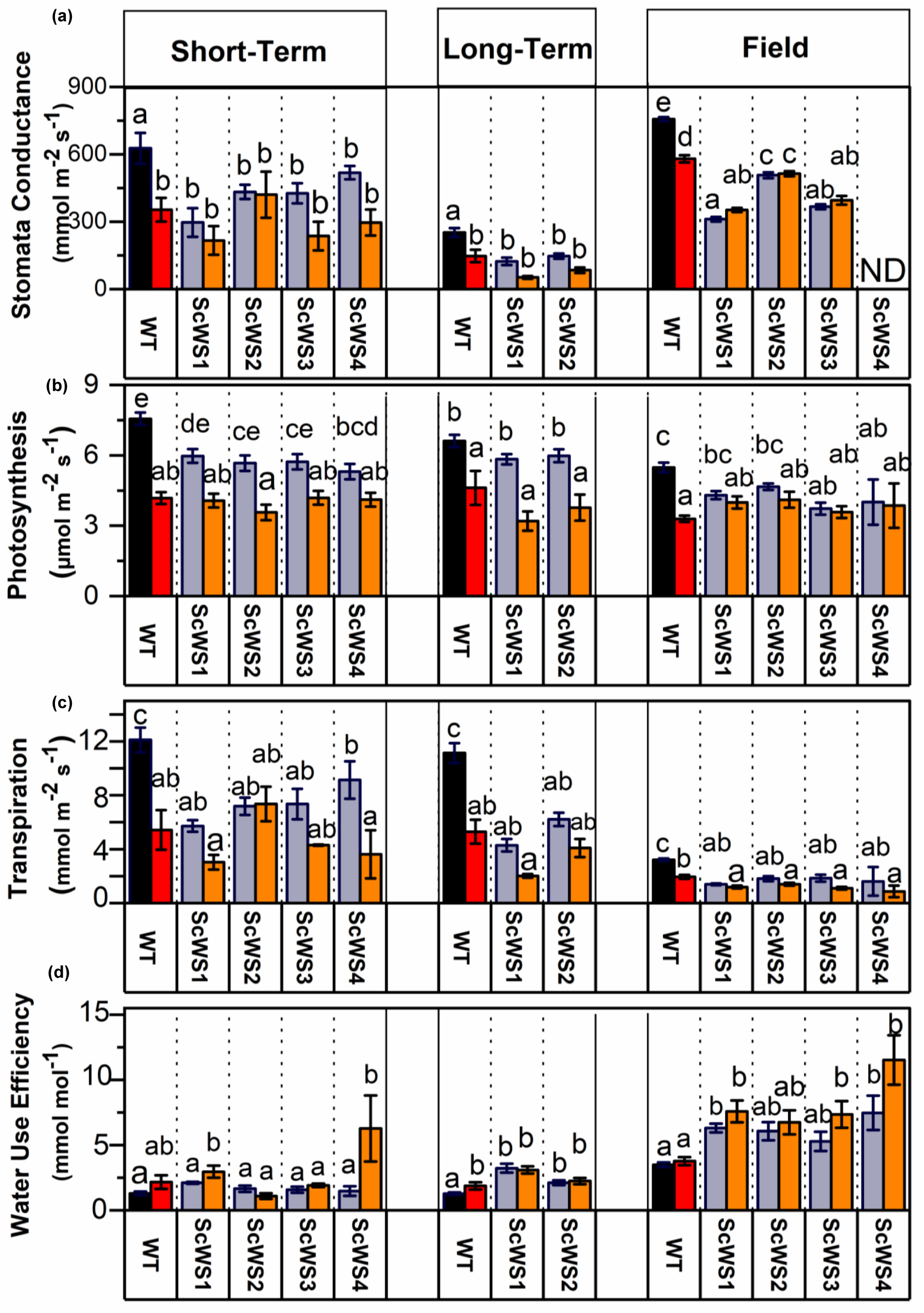
Stomatal conductance (a), photosynthesis (b), transpiration (c) and water use efficiency (WUE) (d) of *P.* x *canescens* (WT) and *ScWS* expressing lines in short-term, long-term and outdoor drought experiments. Gas change measurements were conducted once per week and stomatal conductance every second day from 10:00 to 15:00 during the drought phase. Black and red bars indicate WT under control and drought-stressed conditions and grey and orange bars the *ScWS* lines under control and drought-stressed conditions. Bars indicate means (± SE) with n = 5 per line per treatment. Different letters indicate significant differences at *p* ≤ 0.05 (ANOVA, *post-hoc* Tukey test).

Although the stomatal conductance in the *Sc*WS lines was about twice lower than in the WT under well-irrigated conditions, photosynthesis was not significantly affected (mean decrease: -15%, Fig. 4b). Under drought, photosynthesis decreased more strongly in the WT than in *Sc*WS lines, reaching similar levels across WT and transgenic poplars (Fig. 4b). To gain deeper insight into the photosynthetic behavior of *Sc*WS lines, we measured CO_2_ assimilation under increasing light intensities. We found that the *Sc*WS lines and the WT exhibited similar photosynthesis rates under low light intensities but higher photosynthesis in the WT than in the *Sc*WS lines under high light (≥ 400 µmol m^-2^ s^-1^ photosynthetically active radiation [PAR], Supplement Fig. S11). This finding indicates that light-limited photosynthesis was not additionally affected by reduced stomatal conductance of the *Sc*WS lines. Nighttime stomatal conductance and transpiration were low and did not differ between the *Sc*WS lines and the WT neither under greenhouse nor under field conditions (Supplement Table S1, Table S2). Thus, the regulation of stomatal closure was not hindered by the occluded stomatal phenotype of the *Sc*WS lines.

In accordance with the greatest stomatal conductance, well-irrigated WT plants also showed the highest transpiration rates (Fig. 4c) and lowest instantaneous water use efficiencies (Fig. 4d). Under drought, WT plants showed either no or small increases (+20%) in the instantaneous water use efficiencies (Fig. 4d). With the exception of short-term experiments, *Sc*WS poplars exhibited higher instantaneous water use efficiencies than the WT poplars (Fig. 4d). The greatest instantaneous water use efficiencies were observed for *Sc*WS poplars during drought stress under outdoor conditions (Fig. 4d).

To further characterize the impact of Sc*WS* expression on water consumption, we determined soil moisture, predawn water potentials, and whole-plant leaf area as decisive factors for plant water balance. When the plants were subjected to long-term drought under greenhouse conditions, the WT plants showed a faster decrease in soil moisture than the *Sc*WS lines (Fig. 5a), indicating higher water consumption per WT plant. Higher water spending of the WT poplars further resulted in a stronger decline in the predawn water potential than in the *Sc*WS poplars (Fig. 5b). Since whole-plant leaf areas (Fig. 5c), the plant stem heights (mean height WT: 95.5 ± 1.4 cm and *Sc*WS lines: 87.7 ± 1.9 cm) and non-stomatal water loss (Fig. 2f) did not differ between WT and *Sc*WS lines, the water-saving phenotype of the *Sc*WS lines can be attributed to partial stomatal occlusion. After three weeks of drought under greenhouse conditions, the WT poplars contained about 25% less leaf area (Fig. 2f) and 20% less leaf biomass (7.8 ± 0.2 g) than the *Sc*WS lines (9.7 ± 0.3, p = 0.035, Kruskal Wallis test), mainly because of greater leaf shedding, whereas stem biomass was unaffected (Fig. 5d).

**Figure 5.**
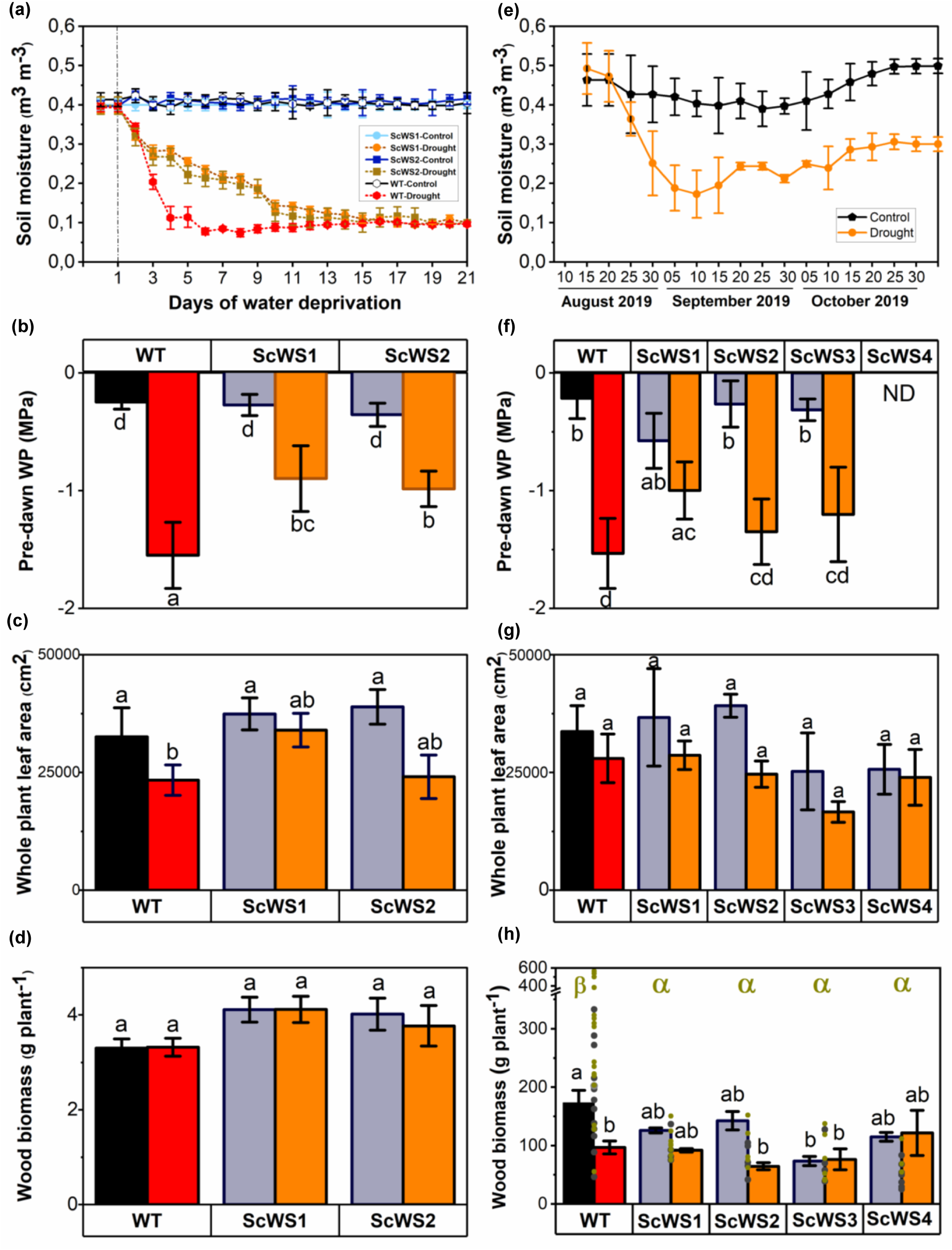
Soil moisture (a,e), pre-dawn water potential (b,f), whole-plant leaf area (c,g) and wood biomass (d,h) of *P.* x *canescens* wild type (WT) and *ScWS* expressing lines in a long-term greenhouse and an outdoor drought experiment. Soil moisture was measured regulary. The predawn water potential was measured at night before dawn using plants 12 d (greenhouse experiment) and 8 weeks after withholding water (outdoor experiment). Data show means (± SE) (n = 8 per line and treatment). Whole-plant leaf area and and wood biomass (stem plus branches) were determined at the time point of harvest (means ± SE of n = 6 individuals in the greenhouse experiment and 5 to 10 in the outdoor experiment 2019) per line and treatment. Black and red bars indicate WT under control and drought-stressed conditions and grey and orange bars the *ScWS* lines under control and drought-stressed conditions. Different letters above the bars indicate significant differences at *p* ≤ 0.05 (ANOVA of log-transformed data, post-hoc Tukey test). In panel (h), bars indicate mean biomass in the first year (2019) and symbols biomass of individual plants in the second year (2020). In 2020, all plants were well irrigated (dark grey symbols = well irrigated in 2019, dark yellow symbols = drought-stressed in preceding year with the exception of wood biomass under outdoor conditions. Transformation did not result in homogeneity of variances. Therefore, the data were analysed by the non-parametric Kruskal Wallis test. Different greek letters in (h) indicate differences at *p* < 0.05 in 2020.

In contrast to plants in individual pots, the outdoor plants were grown in mixtures of WT and *Sc*WS lines, sharing large soil volumes (Supplement Fig. 10). The water consumption of individual plants could therefore not be assessed. Withholding water resulted in slow changes in soil moisture (Fig. 5e) and caused similar reductions in the pre-dawn water potentials of leaves from WT and *Sc*WS lines (Fig. 5f). *ScWS* expression of the field-grown plants was three orders of magnitude higher than that of *PcWDS1* in the WT and showed marginal increases under drought (Supplement Fig. 12). *Sc*WS and WT had similar leaf areas, which were not significantly affected by drought (Fig. 5g).

Since plant fitness is also affected by herbivory, we scored the presence of herbivores (caterpillars and leaf-feeding beetles) on the leaves and estimated their relationship with the scores for leaf damage and predatory insects (i.e. beneficial insects such as ladybugs and wasps) at four time points. Poplar genotype, position of the plants at the edge or in the middle of the plantations, and drought treatment were also included as environmental factors, potentially affecting herbivory. Plant genotype (WT or *Sc*WS lines) and the position were unrelated to herbivores and therefore, removed from the model (Table 1). Leaf damage, predatory insects, time point, and drought treatment were significant factors explaining the variation of herbivores on leaves (Table 1).

**Table 1:** Estimated regression model for the relationship of herbivorous insects with leaf damage, predatory insects, planting pattern and drought. Scores (scale from 1 to 5) for insects and damage were determined four times (mid-August, end of August, mid-September and end of September) on drought-stressed and well-irrrigated poplars (= treatment). The factors position (edge or inside of the mini-plantation) and genotype (WT, *ScWS1*, *ScWS2*, *ScWS3*, *ScWS4*) were not significant and removed from the model. Data show the results of a negative binomial regression model. Percentage of deviance explained by model = 33.2% and adjusted percentage = 25.8%

| Source | Deviance | Df | *P*-Value |
| --- | --- | --- | --- |
| Model | 63.37 | 6 | <0.0001 |
| Residual | 127.70 | 313 | 1.00 |
| Total (corr.) | 191.07 | 319 |  |

| Likelihood Ratio Tests |  |  |  |
| --- | --- | --- | --- |
| Factor | Chi-Square | Df | *P*-Value |
| Leaf damage | 4.67 | 1 | 0.0307 |
| Predatory insects | 8.43 | 1 | 0.0037 |
| Treatment | 16.71 | 1 | <0.0001 |
| Time | 11.30 | 3 | 0.0102 |

At the end of the field season, the aboveground plant biomass was harvested. Since the position of the plants (inside or at the edge of the plantation) did not affect biomass production (p = 0.294, Kruskal Wallis test), all plants were included in the analyses. Well-irrigated WT produced the highest stem biomass and line *Sc*WS3 the lowest, while the *Sc*WS lines 1, 2, and 4 had an intermediate stem biomass (bars in Fig. 5h). Across all treatments, drought caused a decline in wood biomass (p = 0.0003, Kruskal Wallis test). However, for a given genotype, only the WT biomass was significantly reduced, while none of the *Sc*WS lines showed significant drought-induced reductions (Fig. 5h).

In short-rotation plantation practice, the ability of stumps to produce new shoots after aboveground harvest is an important feature. In the following growth season, we observed the re-sprouting of new shoots for 87% of the *Sc*WS lines and for all WT plants. During the growth season, gas exchange of the *Sc*WS lines and the WT was similar to that in the previous season, however, with a stronger restriction of photosynthesis (Supplement Table S3). At the end of the growth season, the woody biomass of some *Sc*WS lines was approximately up to 65% lower than that of the WT (individual data points in Fig. 5h). Previously drought-stressed plants did not show significantly different woody biomass compared with well-irrigated plants (p = 0.061, Kruskal Wallis test).

## Discussion

### Sc*WS* expression enriches the total wax content in poplar leaves and results in aberrant wax accumulation at the stomatal ledges

In this study, we employed *P*. x *canescens*, an important, highly water-spending biomass species (Xi *et al*., 2021) to test whether the heterologous expression of a biotechnologically relevant enzyme, the *Sc*WS can improve cuticular properties and reduce water consumption. This question is important because poplar plantations are known for their huge water demand, which may even cause a decline in the groundwater table (Wilske *et al*., 2009; Folch and Ferrer, 2015). In previous studies, the focus of transgenic Sc*WS* expression was on increasing wax ester concentrations in seeds using model plants (*A. thaliana*) and crop plants such as *C. sativa* (Yu *et al*., 2018; Iven *et al*., 2016). Significant enrichment of wax esters in *A. thaliana* and *C. sativa* required co-transformation with FAR enzymes suggesting that the production of wax esters in seeds was limited by the availability of precursors (Iven *et al*., 2016; Vollheyde *et al*., 2021). Enhanced production of wax esters in leaves of *Nicotiana benthamiana* also required *FAR* coexpression (Domergue and Miklaszewska, 2022). Our study shows that in poplar *FAR/*Sc*WS* coexpression was not necessary to enhance the wax ester content, at least to some extent, since the leaves of the *Sc*WS lines contained increased lipid droplets and enhanced wax ester levels, despite strongly decreased expression levels of Pc*CER4/FAR3* compared with the WT. The Sc*WS* expression caused down-regulation of transcripts in both the alcohol- and the alkane-forming pathways (i.e., *WSD4, CER4/FAR3, CER1*) and resulted in decreased amounts of disctinct products of these pathways (primary alcohol, alkanes, wax esters) in the cuticle. Therefore, we speculate that the intracellular accumulation of wax esters may cause feedback inhibition of wax biosynthesis at least in epidermal and mesophyll cells, possibly already close to the entrance of the pathway (Fig. 6). Another possibility would be that coordinated regulation co-limits the two branches of wax biosynthesis (Yang *et al*., 2022). These suggestions, especially with regard to the proposed function of *WSD4* in wax ester biosynthesis, require further investigations.

**Figure 6.**
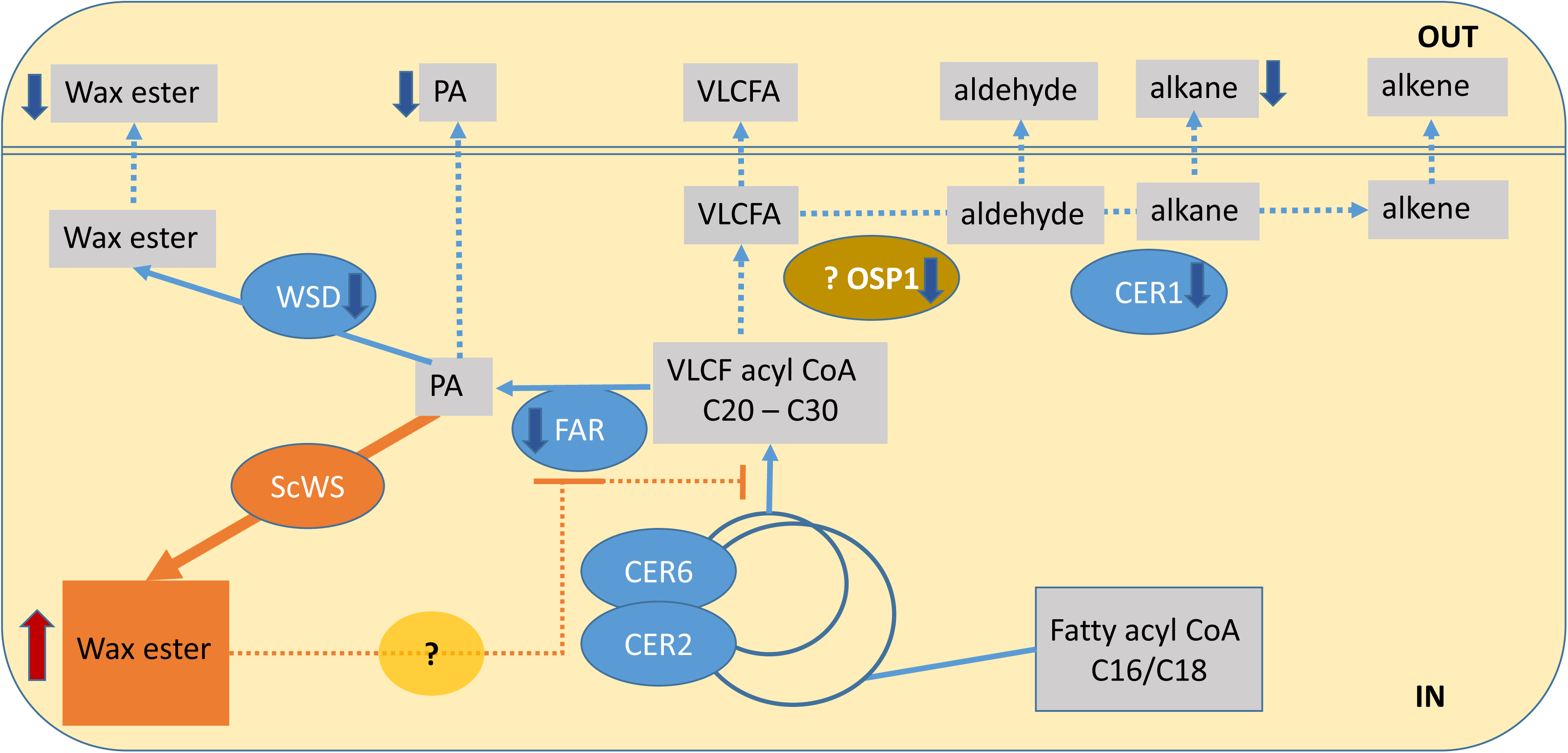
Proposed mechanism of wax biosynthesis in the jojoba wax ester synthase (*ScWS*) overexpressing poplar lines. *Sc*WS activity results in increased intracellular wax esters. To control carbon flux, the biosynthetic pathway for wax esters is down-regulated involving CER4/FAR3 (FAR) and WSDs (here WDS4) but not WDS1. As the consequence of decreased FAR, PA (primary alcohols) decrease. In the alkane forming pathway, *CER1* and alkanes were decreased. *OSP1* expression was localized specifically in the guard cells of stomata and positioned upstream of *CER1* (Tang et al. 2020). OSP1 may convert VLCF acyl CoA into VLCA (= very long chain fatty acid) as precursors for the alkane pathway as suggested by Tang et al. (2020). The suppression of OSP1 may deregulate the localization VLCF aliphatic compunds, causing the occluded stomatal phenotype. The collective suppression of key genes of wax biosynthesis suggests a feed-back inhibition close to the entrance of the pathway by an unknown mechanism.

The observed accumulation of lipid droplets and wax esters was an important result confirming that the *Sc*WS enzyme was active *in planta*. However, we did not observe increased cuticular wax esters or increases in any other substance classes that typically constitute total wax load (Samuels *et al*., 2008; Kunst and Samuels, 2009; Dimopoulos *et al*., 2020). Grünhofer *et al*. (2024) studied the surface waxes of *P*. x *canescens* in a CRISPR/Cas9 *cer6* mutant. In the light of our study, one might have expected that lack of CER6, which is a key enzyme at the entry of wax biosynthesis, may restrict the production of cuticular waxes. On the contrary, the *cer6* mutant showed increased C44 wax esters and enhanced non-stomatal water loss (Grünhofer *et al*., 2024). This unexpected result can be explained by our observations that increased intracellular wax esters may mediate reduced cuticular water loss. Grünhofer *et al*. (2024) speculated that enhanced C44 wax esters might result in structural disturbance of the surface by forming more pores that allow unspecific water loss. However, the reasons for their finding remain enigmatic.

In our study, we found increased expression levels of *CER6, CER2, and WSD1* under drought stress in WT leaves but no influence on the total wax load, its composition or on non-stomatal water loss. Altogether, these findings support that WT leaves regulate wax biosynthesis to maintain functional homeostasis under drought stress. This was apparently also true for *Sc*WS lines since non-stomatal water loss was neither compromised nor enhanced compared with WT plants. Obviously, the expression of Sc*WS* in poplar could not reinforce the cuticle to an extent achieved for example by overexpression of *WDS1* in *A. thaliana* and *C. sativa* (Abdullah *et al*., 2021). Our results support that maintenance of cuticular tightness in drought-stressed *Sc*WS lines was achieved by increases in C27 alkanes and C28 primary alcohols, compensating for the reductions of these compounds present in non-stressed *Sc*WS poplars. In conclusion, our hypothesis that overexpression of Sc*WS* can reduce cuticular water loss was not supported. Instead, we found accumulation of intracellular lipids, which may be interesting for biotechnological applications. This potential of our novel *Sc*WS poplar lines deserves further attention but is beyond the scope of this study.

### Sc*WS* expression shows an occluded stomatal phenotype and improved water use efficiency through an aberrant wax accumulation at the stomatal ledges

An important unexpected result of our study was that *ScWS* expression in poplar phenocopied the occluded stomata syndrome. Tang *et al*. (2020) found that the *A. thaliana osp1* null mutation causes partially occluded stomata, which have a similar phenotype as those found here in *ScWS* expressing poplars. In our study, the *ScWS* expression caused a drastic suppression of *PcOSP1*, suggesting a lack of OSP1 might have been the cause of occluded stomata. Whether the *ScWS* under the constitutive *pS35* shows different local activities in epidermal and guard cells with the observed downstream consequences for *OSP1* expression and stomatal development, remains to be elucidated. Tang *et al*. (2020) proposed that OSP1 (a GDSL lipase) may be a thioesterase since it can hydrolyze C22:0 and C26:0 acyl-CoAs *in vitro* (Tang *et al*., 2020). *AtOSP1* was localized to the guard cells and appeared to be part of the alkane forming pathway, acting upstream of CER1 (Tang *et al*., 2020). The exact molecular mechanisms *in planta* that result in stomatal occlusion are still unknown but point in concert with our study and other investigations (Gray *et al*., 2000; Hunt *et al*., 2017; Tang *et al*., 2020) to a central role of wax biosynthesis as a hub controlling cuticle enforcement jointly with proper stomatal development.

Stomatal pores form between a pair of guard cells by a series of developmental steps, involving localized cell wall thickening and separation to create the pore (Willmer and Fricker, 1996). At the upper side of the ventral wall around the pore, cuticle waxes extend forming “lips” or so-called outer cuticle ledges (Hunt *et al*., 2017). The formation of the ledges is controlled by FOCL1 (FUSED OUTER CUTICLE LEDGES 1) (Hunt *et al*., 2017) and OSP1 (Tang *et al*., 2020). Null mutants of these genes independently cause a high fraction of occluded stomata. The *osp*1 mutant has a reduced cuticular wax load, exhibits increased non-stomatal evaporation but decreased stomatal transpiration (Tang *et al*., 2020). In our study, non-stomatal water loss from leaves and night-time respiration did not show differences among the transgenic *Sc*WS lines and the WT, supporting that the moderate decrease in the total cuticular wax load observed in the transgenic poplar *Sc*WS lines did not influence the cuticular barrier properties for water. This finding agrees with previous studies reporting that not the amounts of cuticular waxes but their composition were decisive to prevent uncontrolled water loss (Schreiber and Riederer, 1996; Riederer and Schreiber, 2001; Meng *et al*., 2019; Seufert *et al*., 2022; Grünhofer *et al*., 2024). Our study shows partial occlusion of stomata by wax esters in the *ScWS* lines. We cannot exclude that the aberrant production of waxes also covers the surface of guard cells but the overall area of guard cells per leaf was obviously too small to affect the total cuticular wax load.

Here, we demonstrate that *Sc*WS expression causes significant increases in instantaneous water use efficiency, which result in decreased whole-plant water consumption and delayed leaf shedding under greenhouse drought conditions. Since no significant influence on the non-stomatal water loss was found, the improved water economy of the *Sc*WS lines was caused by partially occluded stomata. In accordance with transgenic *A. thaliana* plants with occluded stomata (Hunt *et al*., 2017; Tang *et al*., 2020), the almost two-fold reduction in stomatal conductance of the transgenic *Sc*WS poplar lines had initially only little or no negative effects on photosynthesis or growth. However, light curves of photosynthesis showed that stomatal limitations of carbon assimilation occurred mainly at light saturation. This finding implies that optimum photosynthetic conditions cannot be exploited in field plantations. Yet, under cloudy weather conditions or shading, the *Sc*WS lines would have no disadvantage with regard to carbon assimilation, while consuming less water. Other factors, which hamper productivity such as infestation with insects and leaf damage were similar in the *Sc*WS lines compared to the WT. Therefore, we conclude that the growth reductions in the second year were the cumulative result of moderate photosynthetic reductions of *Sc*WS lines under field conditions. However, the improved water use efficiency is expected to be of a huge benefit for plantations which are currently irrigated or tap on ground water, causing problems for sustainable management (Wilske *et al*., 2009; Xi *et al*., 2021).

In conclusion, our study provides evidence that bioengineering of the stomatal ledges opens a promising avenue to improve water management of plantations in a changing, drier climate. However, towards this goal, a deeper understanding of the molecular mechanisms is required to enable fine-tuning of dedicated genes. To date, only three independent studies (Hunt *et al*., 2017; Tang *et al*., 2020; our study), each targeted at a different enzyme, show that increased water use efficiency can be achieved by increased wax accumulation at the stomatal ledges. Our study provides additional insights revealing that poplars with this syndrome have stable *Sc*WS expression, reasonable fitness but reduced productivity under field conditions. An obvious next step is a better control of stomatal wax accumulation, for example, under a drought-inducible promoter to optimize the balance between water use efficiency and growth.

## Materials and Methods

### Poplar cultivation and transformation

Stock cultures of *Populus* x *canescens*, a hybrid of *P*. *alba* x *P*. *tremula* (clone INRA 717-1B4) were micropropagated under sterile conditions (Müller *et al*., 2013) and used for Agrobacterium-mediated shoot transformation according to protocols of Bruegmann *et al*. (2019) and Muhr *et al*. (2016). The details have been reported in Supplementary Methods S1. Regenerating plantlets from independent transformation events were tested for the presence of the target gene *ScWS* by Sanger sequencing (Microsynth Seqlab, Göttingen, Germany). Successfully transformed lines carrying the *p35S::ScWS* construct were denominated *ScWS1*, *ScWS2*… *ScWSn*. For the experiments we used *ScWS1* to *ScWS4*.

### Short-term, long-term, and outdoor growth phenotyping and drought experiments

WT and *ScWS* lines from sterile stock cultures were multiplied by stem microcuttings and cultured on ½ MS medium (Murashige and Skoog, 1962) with a 16 h light / 8 h dark photoperiod until the plantlets were well rooted. Approximately six-week-old plantlets were directly planted into soil [two parts (*v/v*) Fruhstorfer Erde Type N, Hawite Gruppe GmBH, Vechta, Germany), eight parts coarse sand (Ø 0.71–1.25 mm; Melo, Göttingen, Germany), and two parts fine sand (Ø 0.4–0.8 mm; Melo, Göttingen, Germany)] and acclimated for about two weeks to greenhouse conditions. For this purpose, the plants were initially covered by a beaker, which was gradually removed (Müller *et al*., 2013). For short-term experiments, 3 L pots and for long-term experiments 7 L pots were used. For the outdoor experiments, a caged area (Göttingen, Germany, 51.55739°N, 9.95857°E, 153m above sea level; Yu *et al*., 2019), with large containers (3m x 3m x 0.5m) filled with a mixture of sand and compost (Vogteier Erdenwerk GmhH, Niederdorla, Germany) was used. Potted plants were acclimated to greenhouse and then to outdoor conditions, before planting them into the outdoor containers. To control water supply, the soil surface was covered by polyethylene foil and the plants were watered by a drip watering system programed and managed by an intelligent controller AQUA PRO Irrigation Controller (NETAFIM, Tel Aviv, Israel).

Short term experiment: We used three cabinets (temperature: 23-24°C, relative humidity ranged from 60-70%). Ambient light was supplemented with additional illumination (3071/ 400 HI-I; Adolf Schuch GmbH, Worms, Germany) to obtain a light period of 16 h and 150 µmol photons m^-2^ s^-1^ PAR at plant height (Yu *et al*., 2019). In each cabinet, 40 plants were randomly distributed. We used 24 plants of the lines *ScWS1*, *ScWS2*, *ScWS3*, *ScWS4* and the WT. The plants were grown for two additional months and used to measure stem height, stem diameter, and leaf number once a week. For the drought treatment, the plants (70 cm height) in each cabinet were divided into two groups, each containing 7 to 8 plants. The control group received 400 mL water per day and the drought-stressed group 100 ml. The soil water content was monitored (HH2 with ML2x sensor, Delta T devices, Cambridge, England) twice a day (8:00 to 8:30h and 15:00 to 15:30h) in the pots of all plants. If the soil water content of a plant fell below the threshold of 0.35 m³ m^-^³ (controls) or 0.09 m³ m^-^³ (severe drought), additional water was administered. After one-week drought treatment, the plants were harvested.

Long term experiment: Poplar WT and lines *ScWS1* and *ScWS2* were grown for three months under the same conditions described for the short-term experiment. Growth was recorded regularly. The plants (n = 8 per line and treatment) were exposed to drought stress by reducing the water supply for three weeks to 100 ml per day instead of 400 mL per day (controls). Then the plants were harvested.

Outdoor experiment: In each container, 49 plants (7 x 7) were planted in October 2018 at a distance of 0.42 m x 0.42 m, resulting in 25 center and 24 border plants. The poplar lines were distributed, so that that all genotypes were mixed (Supplement Fig. S10, Planting scheme). The plants were watered for 10 min twice a day (8:00h and 12:00). In the following year (2019), the plants were initially watered twice a day for 10 min and from end of July until September three-times a day for 30 min and then until the end of the season three times a day for 10 min. Drought-exposed plants were not watered from August to October. Soil humidity was measured weekly (HH2 with ML2x sensor). Air temperatures (°C) were monitored by a weather station next to the experiment. Means per season (data [°C] for year 1/year 2) were: winter 3.5/4.8, spring 9.1/8.8, summer: 18.7/17.9, fall: 10.3/10.4. Stem height and diameter were measured regularly. According to plantation practice, the poplars were set back to the stool at the end of the growth phase in October 2019 by cutting the stem 10 cm aboveground. Leaves and wood biomass were separated, dried at 70°C and weighed. In the following year, resprouting and growth were observed. All plants were watered regularly by the irrigation system. The plants were harvested at the end of the growth season in October 2020.

### Insect and damage scoring of outdoor plants

Scoring was conducted four-times in 2019 (8^th^ and 30^th^ August, 4^th^ and 19^th^ September). We used scores for herbivores (caterpillars [Lepidoptera], poplar leaf beetle [Chrysomela sp.]) from 0 (no herbivores) to 4 (completely covered), leaf area damage from 1 (intact) to 5 (only leaf ribs left) and presence or absence of beneficial insects (ladybugs [Coccinellidae], wasps [Hymenoptera]).

### Harvest

Whole-plant leaf and stem biomass were weighed (fresh and dry). Leaf area was determined by scanning (Image J, https://imagej.net/ImageJ) and weighing samples of fresh leaves and upscaling to whole-plant leaf area by multiplication with whole-plant leaf biomass. Aliquots of fresh tissue (leaf, bark, wood) were immediately shock-frozen in liquid nitrogen and stored at -80°C.

### Gas exchange

Gas exchange (photosynthesis, stomatal conductance, transpiration) was measured on fully expanded, light-exposed leaves, usually leaf number 7 from the stem apex with a multiphase Flash™ Fluorimeter (LI-6800, LI-COR, Lincoln, U.S.A.). Daytime measurements were conducted between 10:00 and 14:00h with 800 µmol photons m^-2^ s^-1^ PAR, leaf temperature of 25-27°C, and ambient CO_2_ of 400 µmol mol⁻¹. Instantaneous water use efficiency was determined as the ratio of net photosynthetic CO_2_ consumption to transpiration (Farquhar *et al*., 1989). Respiration, transpiration and stomatal conductance at night were measured between 02:00 and 06:00h in darkness. The measurements were conducted on n = 5 plants per line and treatment.

### Predawn leaf water potential

Predawn water potentials (SKPM 1400/40 pressure chamber, UP GmbH, Ibbenbüren, Germany) were measured on the petiole of detached leaves (Scholander *et al*., 1964) at night. Leaves from the same position of the stressed and control plants were used.

### Water loss assay

We used fully expanded leaves formed during drought and control leaves of the same age for the water loss assays. Ten leaf disks (Ø 14 mm) were punched out per leaf. The leaf disks floated on distilled water at 23 ± 0.5°C in darkness overnight. The water-saturated leaf disks (SW) were surface-dried, weighed and exposed in an open glass petri dish with the stomatal side on the glass surface. The weight of the disks was determined regularly for 5h (WT_1_, WT_2_…, WT_5_). The leaf disks were oven dried at 80°C for 24 hours and weighed (DW). Relative water loss was determined as:

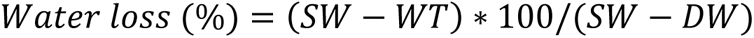

plotted against time, and used to determine the slope of a linear regression from 1 h to 5 h exposure. We did not include the start (0 h to 1h ) because differences in stomatal conductance and unspecific water loss via the injured edges affected water loss in the initial phase. Mean slopes (% h^-1^) were determined for 4 independent biological replicates per treatment and line. Several independent repeated experiments were conducted with similar results.

### Scanning electron microscopy (SEM) of leaf surfaces

Fresh leaves of greenhouse and field-grown WT and *ScWS* poplars were used for SEM analyses as described by Dreischhoff *et al*. (2023).The details have been presented in the Supplementary methods S1. A minimum of three independent biological samples were analysed per treatment and line.

### Transmission Electron Microscopy (TEM) of leaf cross sections

Small leaf segments (max. 0.2 x 0.2 cm) were collected, fixed in glutaraldehyde (Science Services GmbH, München, Germany) overnight and post-fixed in OsO_4_ (Science Services GmbH) at room temperature for 1 h. Then, the samples were dehydrated in an ethanol series and embedded in Araldite-Epon resin according the manufactureŕs instructions (Araldite 502/Embed 812 Kit; Science Services GmbH). Ultrathin cross sections (70 nm) were obtained with an Ultracut E microtome (Reichert-Jung, Heidelberg, Germany) and a diamond knife. The sections were placed on Formvar-coated nickel grits (Plano, Wetzlar, Germany) (Miklaszewska *et al*., 2023). After contrasting with UranyLess (Science Services GmbH) and 3 % lead citrate (Science Services GmbH) following the manufactureŕs instructions, pictures were captured by transmission electron microscopy at 80 kV using a Technai G2 Spirit (FEI company, Eindhoven, Netherlands).

### Confocal laser scanning microscopy (CLSM) of leaf sections

Fresh leaf samples were stained with LipidSpot-488 (Lipid Droplet Stain, Biotium, Fremont, USA) according to manufactureŕs protocol with minor modifications: samples were incubated in darkness at 37°C for 10 min in a solution of 1 µl LipidSpot-488 in 1000 µl phosphate buffer. The stained specimens were embedded on slides with 10 µl Roti embedding flour care (Carl Roth GmbH+Co.KG, Karlsruhe, Germany). Fluorescence was detected by a CLSM (TCS-SP8, Leica, Wetzlar, Germany) with excitation wavelengths of 480-490 nm and emission wavelengths of 580-620 nm through the plane of the leaf (Supplement Fig. 8 d,e,f). Images were recorded at 100x magnification and used to count open, semi-open, and occluded stomata and to count lipid droplets per area unit. Five biological replicates were analyzed per line. Each biological replicate was the mean of two replicates per sample.

### Cuticular wax analysis

For cuticular wax extraction, three leaf discs per plant of 14 mm in diameter with an area of 2.5 cm² were collected, exposed for 30 s to chloroform containing *n*-tetracosane as the internal standard and further processed as previously described (Haslam and Kunst, 2013). Derivatized samples were separated by gas chromatography and mass spectroscopy as described in detail in Supplementary methods S1. The internal standard was used for the quantification of compounds.

### RNA extraction, cDNA synthesis and quantitative real time polymerase chain reaction (qRT-PCR) of genes involved in wax biosynthesis

Frozen tissues were ground in a pre-cooled ball mill (Retsch, Haan, Germany) to a fine powder. The frozen powder (150 mg) was used for total RNA extraction using the CTAB method (Chang *et al*., 1993). Genomic DNA was removed according to the manufacturer’s instructions using the Turbo DNA-free kit by treating with 10 μg DNase (Turbo DNA-free kit, Ambion, Austin, TX) at 37°C for 30 min. The integrity of the purified RNA (2.5 μg) was assessed by agarose gel electrophoresis (Fleige and Pfaffl, 2006). DNase-treated total RNA (5 µg) was used as starting material for double-stranded cDNA synthesis using Oligo(dT)18 primer and RevertAid™ First Strand cDNA Synthesis Kit (Thermo Scientific, Sindelfingen,Germany) according to the manufactureŕs protocol.

Quantitative Real-time PCR (qRT-PCR) was performed in 96-well-plates (qTOWER3 G touch, Analytik Jena, Jena, Germany). Primers for genes of the wax biosynthesis pathway were designed on the basis of *P. trichocarpa* and *P.* x *canescens* sequences (Mader *et al*., 2016) with the Perl Primer software version 1.1.20 (Marshall, 2004) (Supplement Table S4). The primer efficiency was controlled by running a dilution series. The qRT-PCRs were carried out with following conditions: starting temperature 95 °C for 2 min, then 45 cycles of 95 °C for 10 s and 55 °C for 20 s and at the end a melting curve analysis with measurements between 72 and 95 °C. The 2^-ΔΔCt^ method was used to calculate the relative expression of the genes using *PtrPPR_2* and *PtrRpp14* as the reference (Livak and Schmittgen, 2001).

### Statistical analyses

The Software R (version 4.2.2, R Core Team, 2017), Origin 2020 (OriginLab^®^, Northhampton, MA, U.S.A.) and Statgraphics Centurion XVII (Statgraphics Technologies, Inc., The Plains, Virginia, U.S.A) were used for statistical analysis. We tested normal distribution by inspecting residuals and homogeneity of variances by Levene’s test. When the data were not normal distributed, we used log10 transformation to meet the statistical requirements. Data were analysed by ANOVA (line and drought as main factors) and a *post-hoc* test to determine significant differences between means at *p* ≤ 0.05. If normal distribution and variance homogeneity were not achieved, non-parametric methods and Kruskal Wallis test were applied. Count data for insect scores were analysed by a negative binomial regression with genotype, time, drought treatment, and position (border plants, inside plants) as categorical factors and scores for leaf damage and abundance of beneficial insects as quantitative factors. Data are shown as means (± SE). The number individual biological replicates and the statistical tests are indicated in the figure legends.

## Author contributions

AA: conducted experiments, analysed molecular and physiological data, synthetized data, wrote the first draft, GJS: generated transgenic poplars, conducted experiments, analysed data, AK and CH: measured and analysed waxes and phytohormones, FH: measured and analysed SEM data, UL: measured and analyzed TEM data, IF: supervision, fund acquisition, scientific advice, AP: research design, supervision, fund acquisition, scientific advice, manuscript writing. All authors commented on and approved the final version of this manuscript.

## Acknowledgements

We are grateful to M. Fastenrath maintenance of poplar stock cultures, to C. Leibecke for growth phenotyping and insect scoring and to S. Freitag for help with sample preparation for lipid and phytohormones measurements. We thank Prof. A. von Tiedemann, Dr. Birger Koopmann, and Yao Wang (Plant Pathology and Crop Protection, University of Göttingen) for making the confocal laser scanning microcope available.

## Funding statement

We acknowledge funding provided by the IRTG ProTect, GRK2172 (project B2.1, M2.2, Deutsche Forschungsgemeinschaft), the Lichtenberg Research program (Niedersächsisches VW Vorab), and publication funds of the University Göttingen supported by DFG grants to project DEAL.

## Conflict of interest disclosure

The authors declare no conflict of interest.

## Ethics approval statement

Permissions to produce and conduct research with transgenic plants (safety standard S1) conditions have been issued the authorities of Lower Saxony.

## Data availability statement

Data have been deposited in Figshare DOI: 10.6084/m9.figshare.25549702. ScWS poplar lines are available upon reasonable request from A. Amirkhosravi (Forest Botany and Tree Physiology, University of Göttingen).

## Supporting materials

### Supporting figures

**Figure S1.** Phylogenetic analysis of bifunctional wax synthase/diacylglycerol acyl transferases (WSD) in *P. trichocarpa* and *Arabidopsis thaliana*.

**Figure S2.** Expression of *PcWSD1* and *PcWSD4* in different tissues of *P*. x *canescens.*

**Figure S3.** Transmission electron microscopy of cross sections of *P*. x *canescens* leaves of the wild type and *ScWS* lines.

**Figure S4.** Scanning electron microscopy of the adaxial leaf surface of the wild type and ScWS lines of *P*. x *canescens*.

**Figure S5.** Scanning electron microscopy of the abaxial leaf surface of the wild type and *ScWS* lines of *P.* x *canescens*.

**Figure S6.** Scheme for the ABA signalling pathway, relative expression of *MYB96* and *MYB94* and concentrations of abscisic acid in *ScWS* lines and wild-type *P.* x *canescens*.

**Figure S7.** Morphology of stomata of wildtype and transgenic *P*. x *canescens* lines under sterile tissue culture conditions.

**Figure S8.** Fluorescence microscopy of lipid droplets in leaves of *P*. x *canescens* wildtype and *Sc*WS lines.

**Figure S9.** Thin layer chromatographic separation of total lipid extracts from leaves of *P*. x *canescens* wild type and *ScWS* lines.

**Figure S10.** Planting scheme of wild type and *ScWS* lines of *Populus* × *canescens* in mixtures under outdoor conditions.

**Figure S11.** Relative expression levels of *ScWS* in transgenic poplars and of *WSD1* in wild-type poplars under outdoor conditions.

**Figure S12.** Light response curve of photosynthesis of ScWS lines and wild-type *P*. x *canescens*.

### Supporting tables

**Table S1.** Respiration, transpiration and stomatal conductance in darkness of well-irrigated and drought-stressed *ScWS* lines and wild-type *P. x canescens* in a long-term greenhouse experiment.

**Table S2:** Respiration, transpiration and stomatal conductance in darkness of well-irrigated and drought-stressed *ScWS* lines and wildtype *P*. x *canescens* under field conditions.

**Table S3**. Gas exchange of *P. x canescens* wild type and *ScWS* lines under outdoor conditions in 2020 (second growth phase).

**Table S4.** List of the primers and Potri numbers for genes used for the cloning and for expression analyses by qRT PCR.

### Supporting Experimental Procedures

**Supporting Methods S1:** Protocols for poplar transformation, scanning electron microscopy of fresh leaf surfaces and for cuticular wax analysis.

## Notes

### Competing Interest Statement

The authors have declared no competing interest.

