## Supplementary material for "Wax ester synthase overexpression affects stomatal development, water consumption and growth of poplars": Figures_Tables_Methods

**Supporting Figures**
**Figure S1.** Phylogenetic analysis of bifunctional wax synthase/diacylglycerol acyl transferases (WSD) in *P. trichocarpa* and *Arabidopsis thaliana.*
**Figure S2.** Expression of *Pc*WSD1 and *Pc*WSD4 in different tissues of *P*. x *canescens.*
**Figure S3.** Transmission electron microscopy of cross sections of *P*. x *canescens* leaves of the wild type and *Sc*WS lines.
**Figure S4.** Scanning electron microscopy of the adaxial leaf surface of the wild type and *Sc*WS lines of *P*. x *canescens*.
**Figure S5.** Scanning electron microscopy of the abaxial leaf surface of the wild type and ScWS lines of *P.* x *canescens*.
**Figure S6.** Scheme for the ABA signaling pathway, relative expression of MYB96 and MYB94 and concentrations of abscisic acid in ScWS lines and wild type *P.* x *canescens*.
**Figure S7.** Morphology of stomata of wild type and transgenic *P*. x *canescens* lines under sterile tissue culture conditions.
**Figure S8.** Fluorescence microscopy of lipid droplets in leaves of *P*. x *canescens* wild type and *Sc*WS lines.
**Figure S9.** Thin layer chromatographic separation of total lipid extracts from leaves of *P*. x *canescens* wild type and *Sc*WS lines.
**Figure S10.** Planting scheme of wild type and ScWS lines of *Populus* × *canescens* in mixtures under outdoor conditions.
**Figure S11.** Relative expression levels of *ScWS* in transgenic poplars and of *WSD1* in wild-type poplars under outdoor conditions.
**Figure S12.** Light response curve of photosynthesis of *Sc*WS lines and wild-type *P*. x *canescens*

**Supporting tables:**
**Table S1.** Respiration, transpiration and stomatal conductance in darkness of well-irrigated and drought-stressed ScWS lines and wild-type P. x canescens in a long-term greenhouse experiment.
**Table S2:** Respiration, transpiration and stomatal conductance in darkness of well-irrigated and drought-stressed *Sc*WS lines and wild-type *P*. x *canescens* under field conditions.
**Table S3**. Gas exchange of P. x canescens wildtype and ScWS lines under outdoor conditions in 2020 (second growth phase).
**Table S4.** List of the primers and Potri numbers for genes used for the cloning and for expression analyses by qRT PCR.

**Supporting Experimental Procedures.**
**Supporting Methods S1:** Protocols for poplar transformation, scanning electron microscopy of fresh leaf surfaces and cuticular wax analysis.

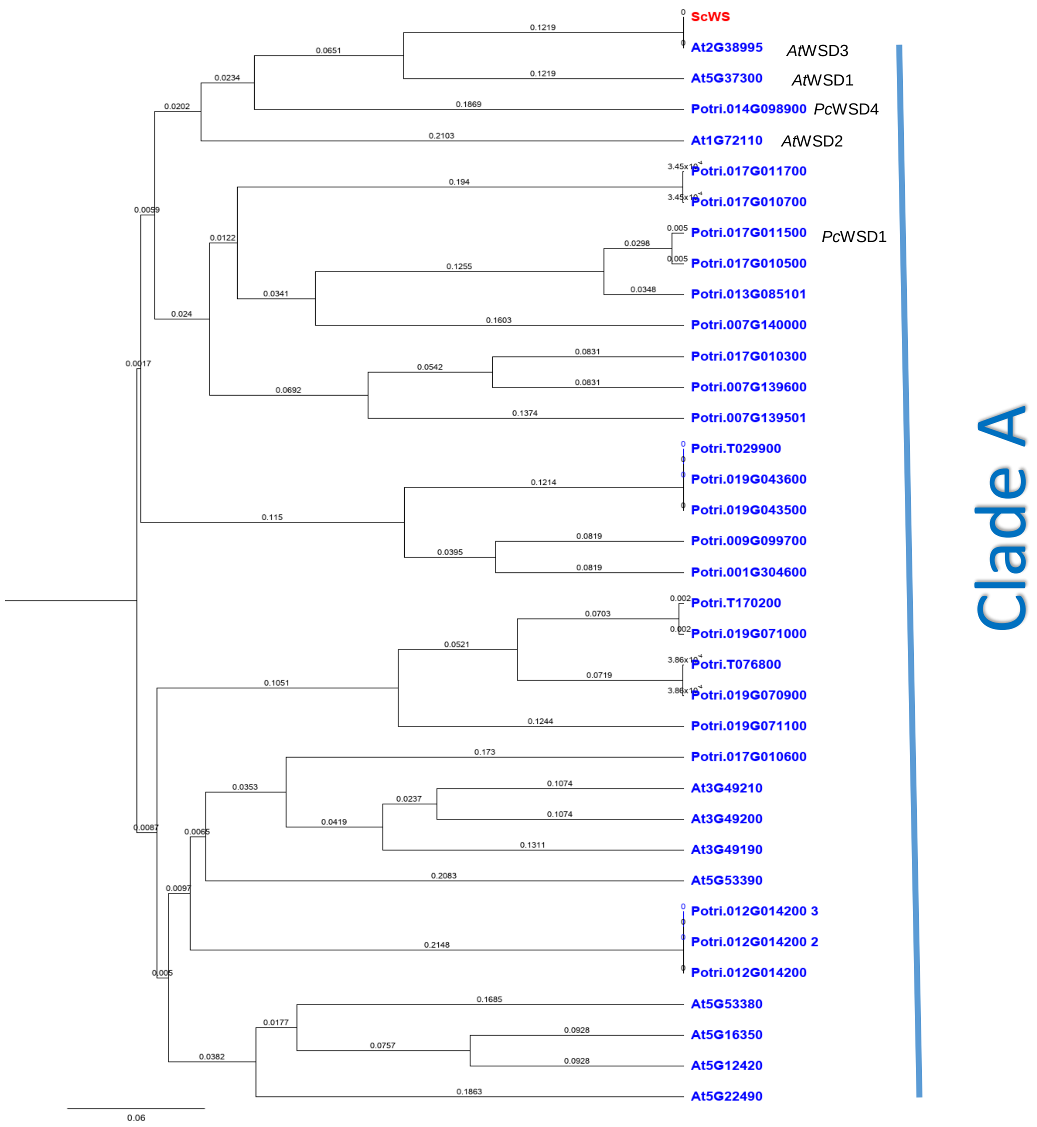


**Supplementary Figure S1: Phylogenetic analysis of bifunctional wax synthase/diacylglycerol acyl transferases (WSD) in *P. trichocarpa* and *Arabidopsis thaliana*.** Clade A was assigned according to Cheng *et al.*, (2022). *A. thaliana* WSD amino acids sequences (AGI numbers from: King *et al.*, 2007) were retrieved from the TAIR database (version 1, accessed on 11-May-2023). The *A. thaliana* WSD amino acid were used to extract the amino acid sequences of the *P. trichocarpa WSDs* from https://plantgenie.org (accessed on 11-May-2023). The phylogenetic tree was built with Geneious Prime (Biomatters, Ltd., Auckland, New Zealand, https://www.geneious.com) by global alignment with free ends gaps alignment type and a cost matrix of 70% similarity (International Union of Biochemistry (IUB) nucleotide ambiguity code, 5.0/-4.5). The genetic distance model Jukes-Cantor with Neighbor-joining tree build method was used with the Jojoba wax ester synthase as the outgroup.

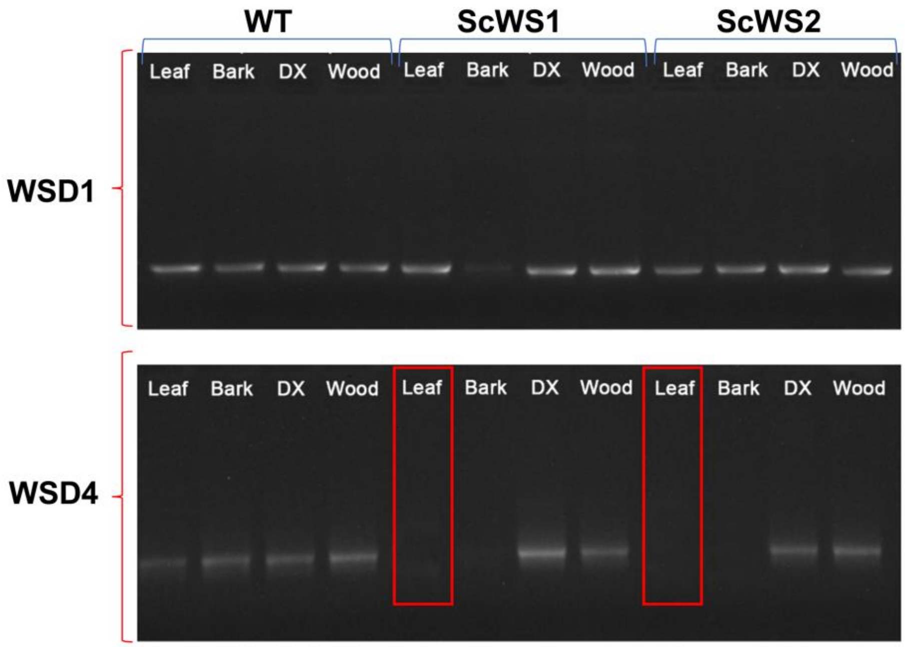


**Supplementary Figure S2: Expression of *Pc*WSD1 and *Pc*WSD4 in different tissues of *P.* x *canescens.*** RNA was extracted from leaves, bark, developing xylem (DX), and wood of wild type (WT) poplar and the lines *Sc*WS1 and*Sc*WS2 lines. RT-PCR was conducted with 10 µl of 70 ng/µl cDNA per slot and the specific primers for *WSD1* and *WSD4* (Supplement Table S3). The products were separated by agarose gel electrophoresis and observed after ethidium bromide staining.

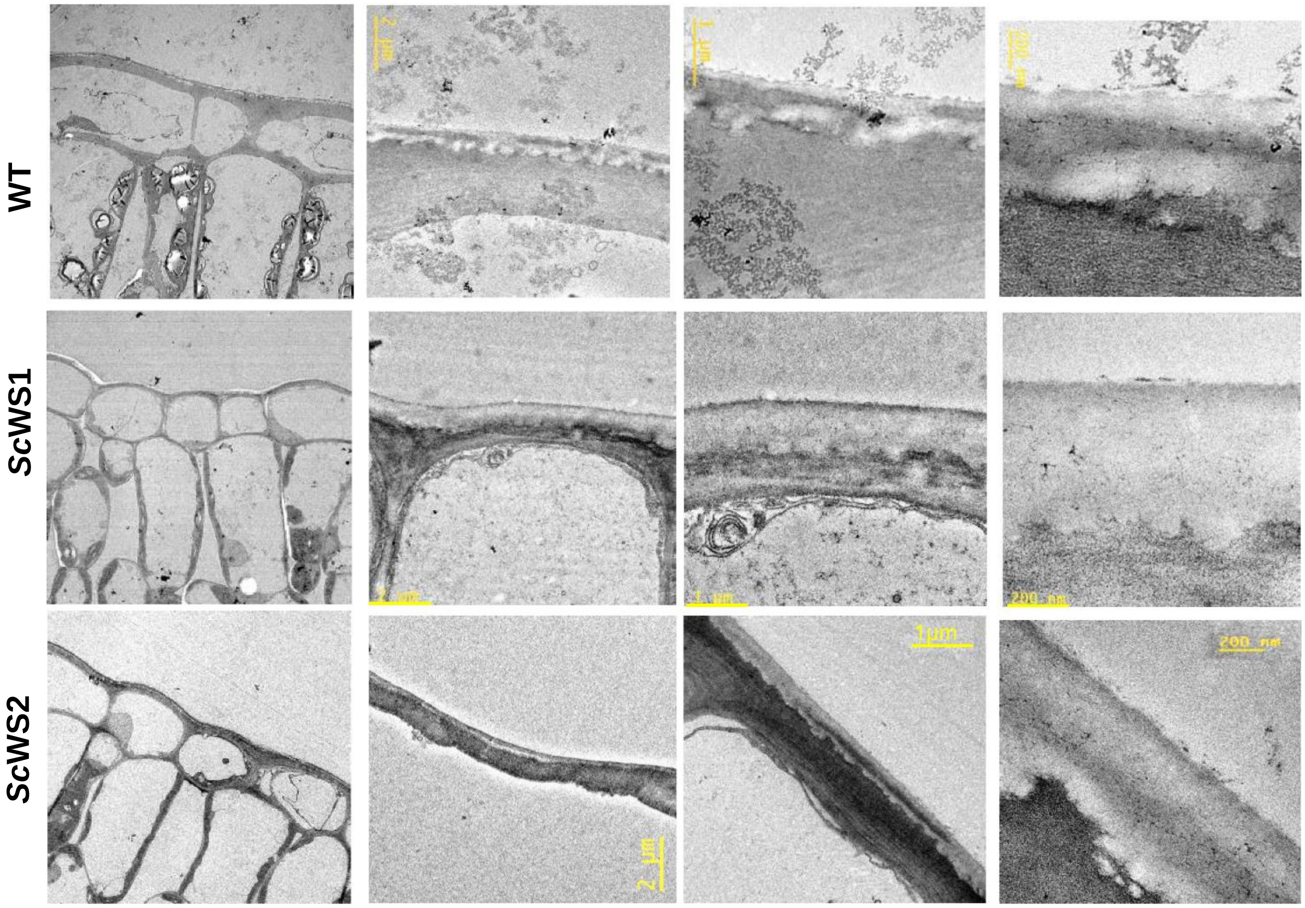


**Supplementary Figure S3. Transmission electron microscopy of cross sections of *P.* x *canescens* leaves of the wildtype and *Sc*WS lines (*Sc*WS1, *Sc*WS2)**. Pictures show the thickness of the cuticle layer with increasing magnification.

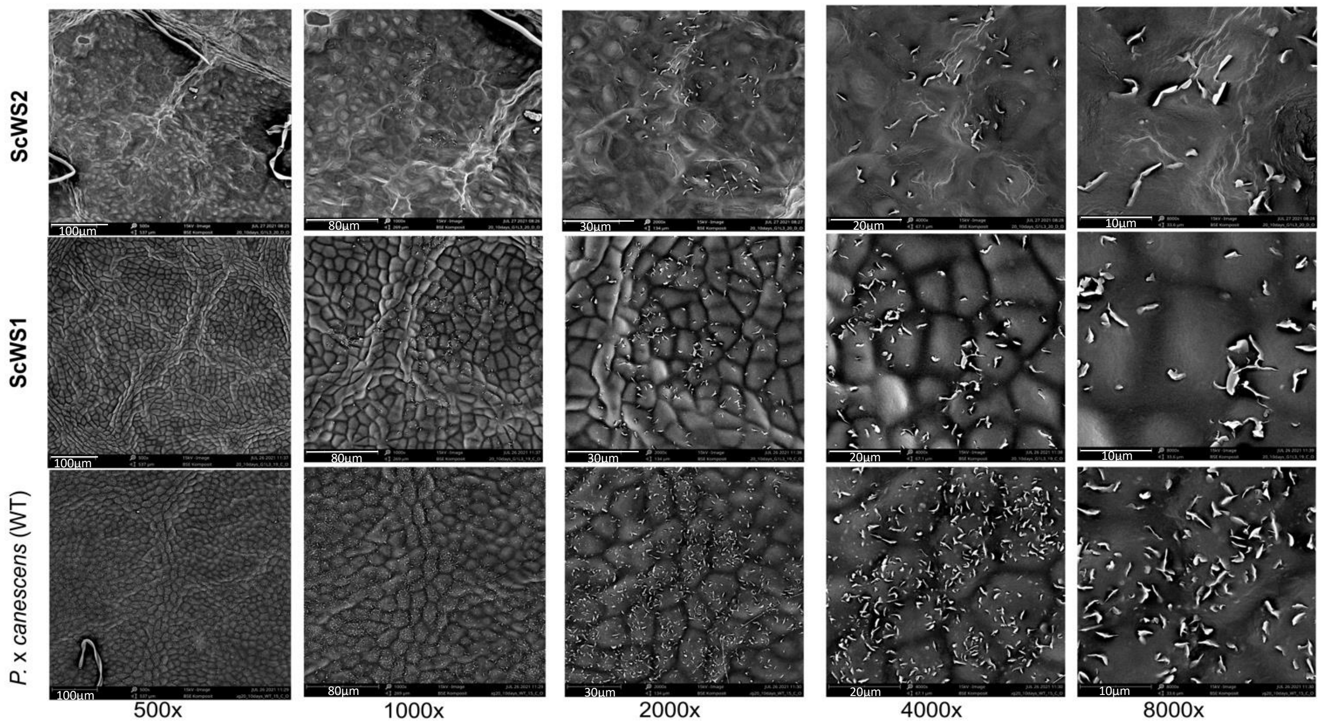


**Supplementary Figure S4**. **Scanning electron microscopy of the adaxial leaf surface of the wildtype and ScWS lines of *P.* x *canescens.*** The pictures are representing the epicuticular crystal wax layer structures at increasing magnifications (from 500X-8000X).

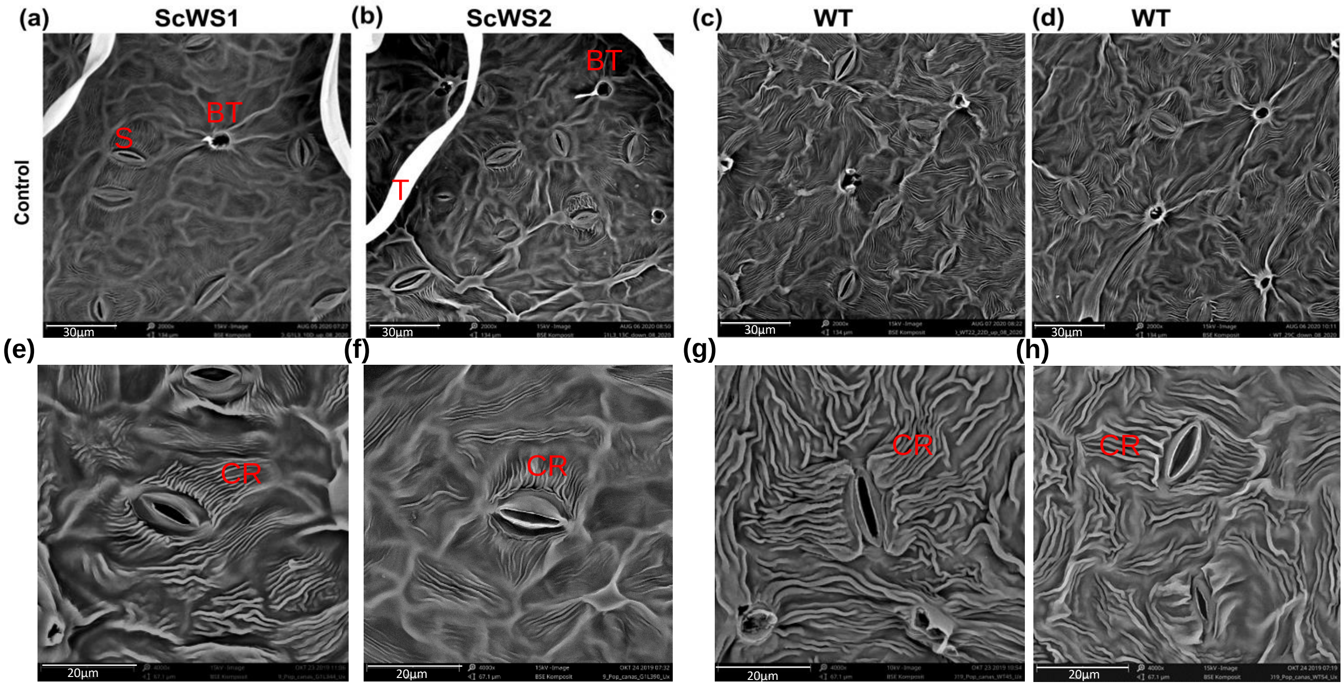


**Supplementary Figure S5**. **Scanning electron microscopy of the abaxial leaf surface of *P. x canescens* *Sc*WS lines** (a,b) **and the wildtype (WT)** (c,d) and the cuticular ridges emerging close to the stomata of the *Sc*WS lines (e,f) and WT (g,h). Abbreviations: (S) Stomata, (BT) Brocken trichome, (T) Trichome, (CR) Cuticular Ridges.

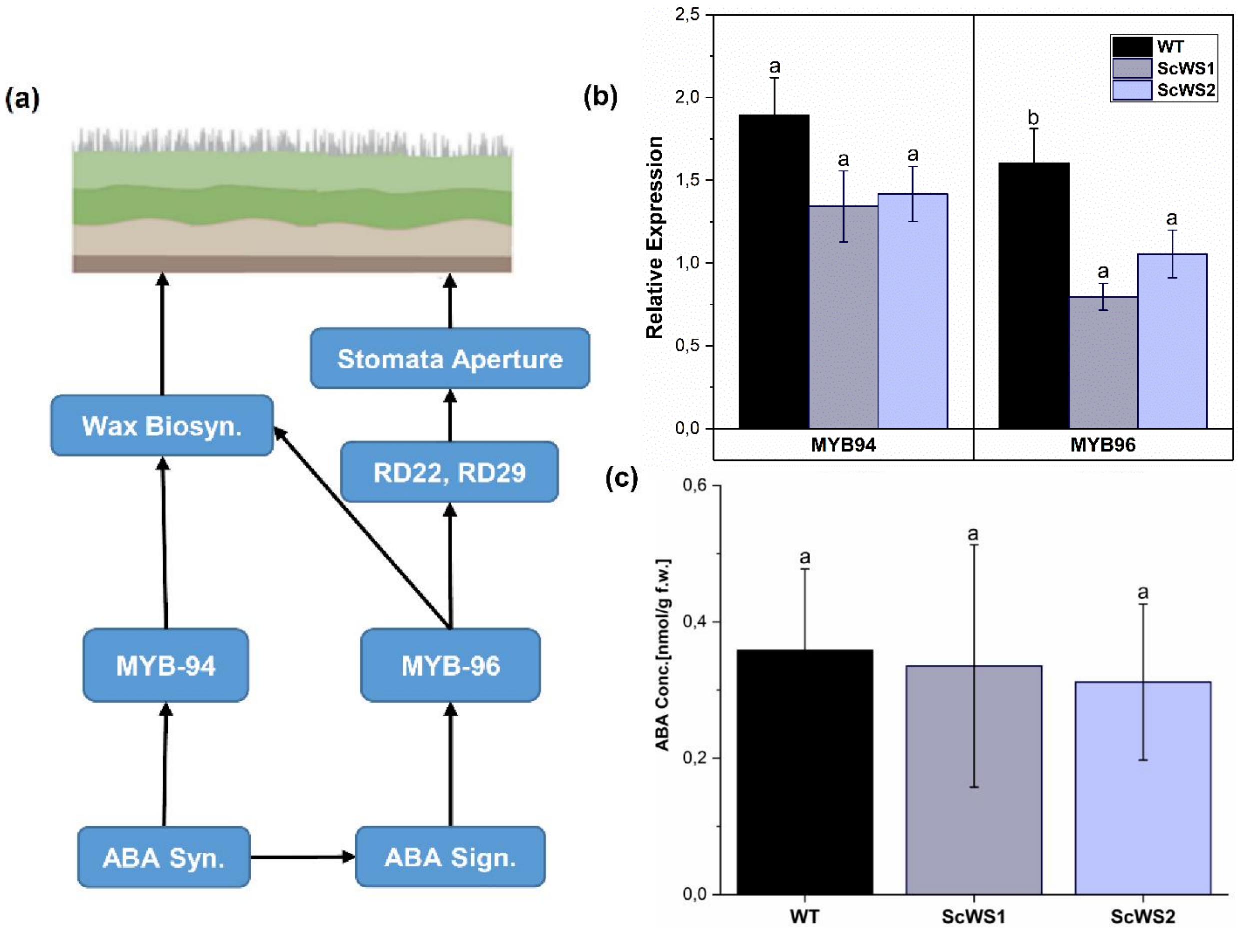


**Supplementary Figure S6**. **Scheme for the ABA signalling pathway in relation to wax biosynthesis and stoma formation (a)**, **relative expression of MYB 96 and MYB 94 (b) and ABA concentrations in leaves of wild type and *Sc*WS expressing *P.* x *canescens* plants**. The scheme was adapted from previous publications (Lee *et al.*, 2020; Lewandowska *et al.*, 2020; Seo and Park, 2010; Seo *et al.*, 2009). Abscisic acid was extracted from frozen leaf powder and determined by liquid chromatography coupled with mass spectroscopy as described by Yu *et al.* (2021). Data show means (± SE, n = 4-7 per line). Different letters indicate significant differences at $p < 0.05$ among the *Sc*WS lines and WT (post hoc Tukey test).

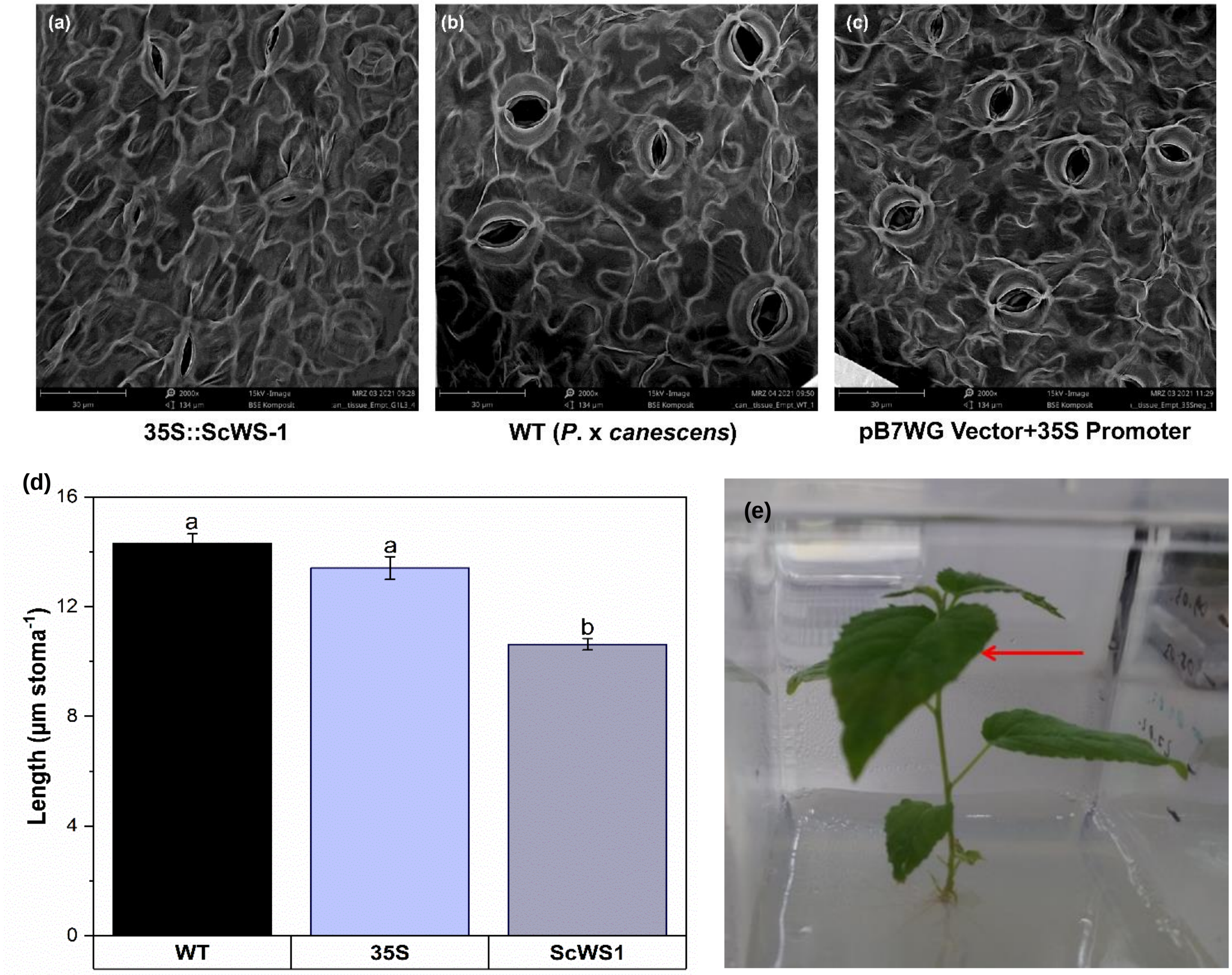


**Supplementary Figure S7. Morphology of stomata and stomatal length of wildtype and transgenic *P.* x *canescens* lines under sterile tissue culture conditions.** Scanning electron microscopy of the abaxial leaf surface (a) line *Sc*WS1, (b) wildtype, (c) empty vector control transformed with the 35S promoter; magnification 2000X. (d) Stomatal length, (e) plants in the tissue culture, red arrow indicates the leaf used for the scanning electron microscopy. Data show means (± SE, n = 4 per line). Different letters indicate significant differences among the lines at P ≤ 0.05 (post hoc Tukey test).

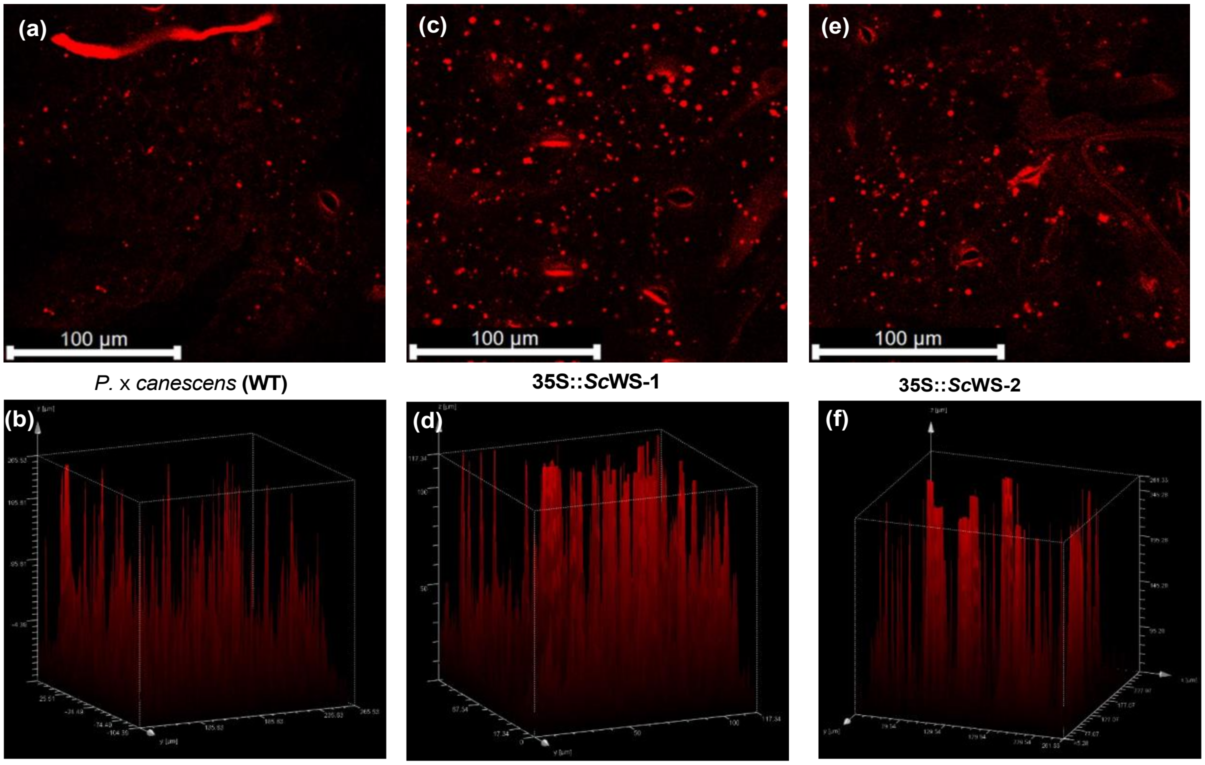


**Supplementary Figure S8. Fluorescence microscopy of lipid droplets in leaves of *P.* x *canescens* wildtype (a,b) and ScWS expression lines ScWS1 (c,d) and ScWS2 (e,f)**. Lipid droplets were visualized by staining with the lipophilic dye Lipid spot II and measured through the leaf plane as indicated by the 3D views (b,d,f).

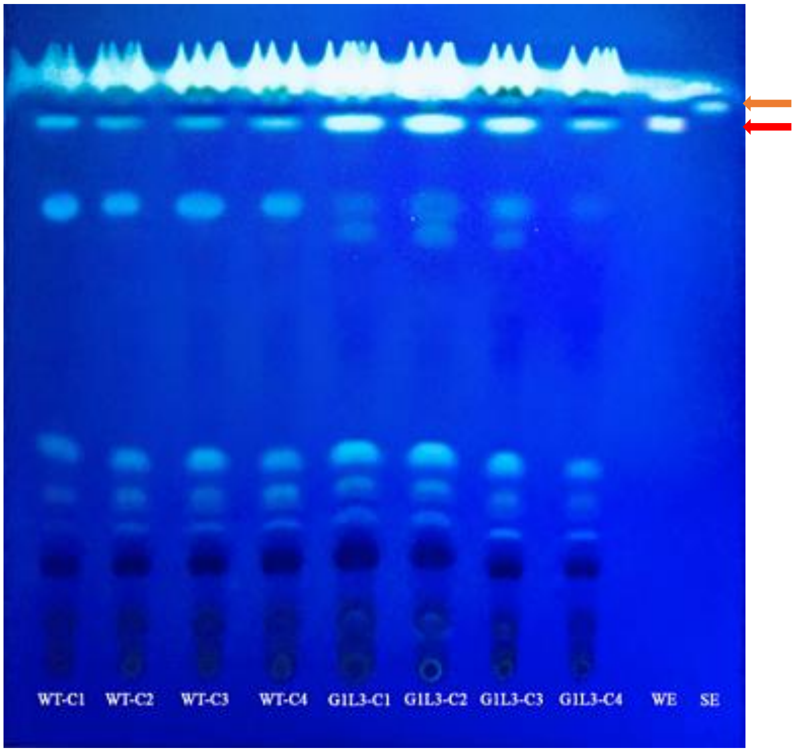


**Supplementary Figure S9. Thin layer chromatographic separation of total lipid extracts of *P.* x *canescens* leaves.** Crude lipid extracts were prepared according to Iven *et al.*, (2013) with minor modifications. Frozen leaf samples (-80°C) were milled and 100 mg were used for extraction in chloroform: methanol (1:1, V/V). The lipid extracts were spotted on 0.25 × 20 × 20 cm F60 silica gel glass plates (Merck, Darmstadt, Germany) and separated by a mobile phase of hexane :diethyl ether : acetic acid (80 : 20 : 1). Wax ester (WE) (red arrow) and sterol ester (SE) (orange arrow) were used as the reference compounds The plate shows an example of several independent extractions. The plates were sprayed with primuline dissolved in 80% acetone and viewed under UV light. WT-C1 to WT-C4 extracts from four individual wildtype plants, G1L3-C1 to G1L3-C4: extracts from four individual *Sc*WS1 plants.

(a)

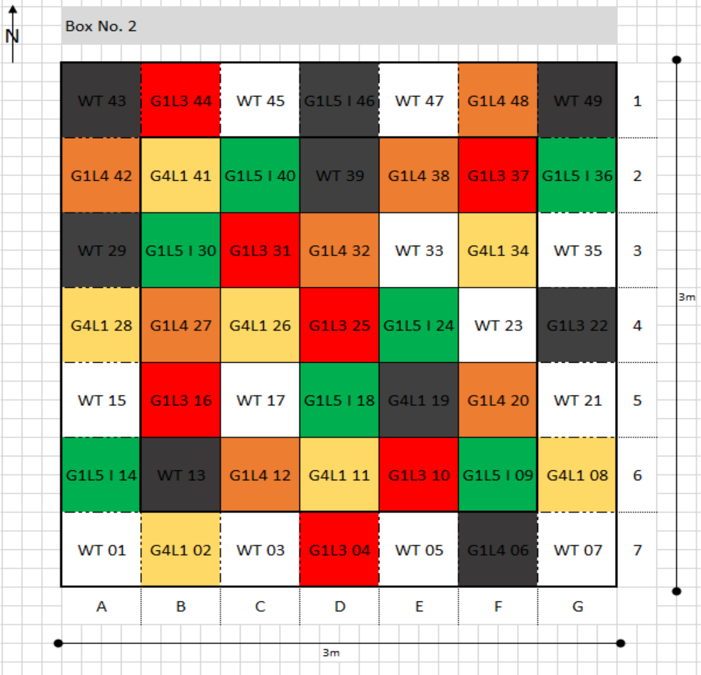


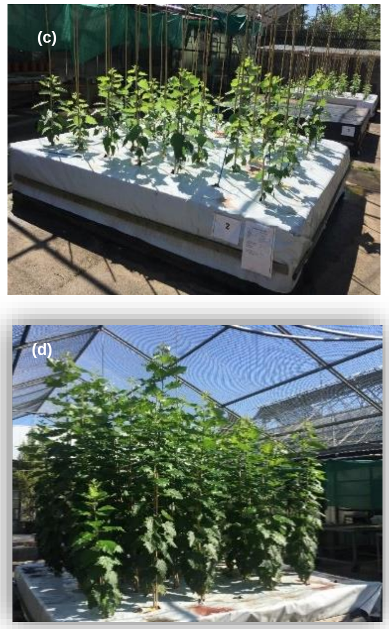


(d)

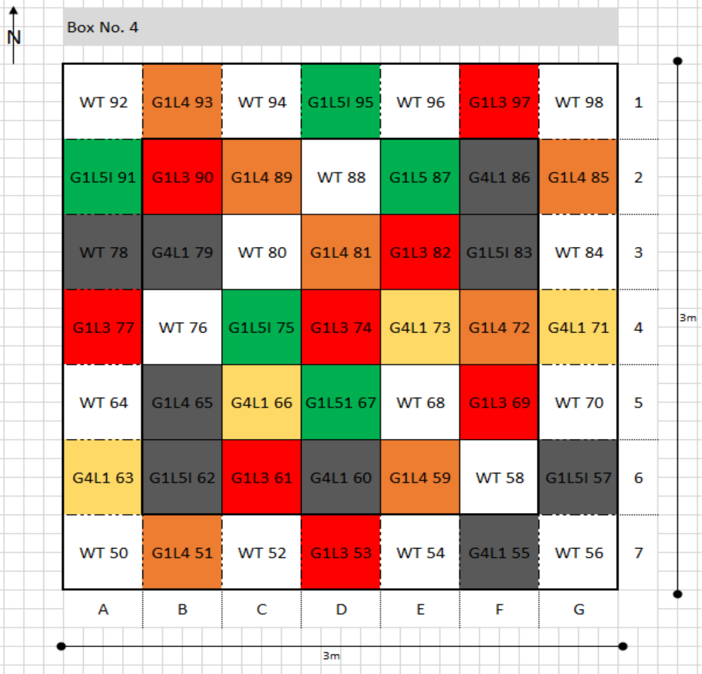


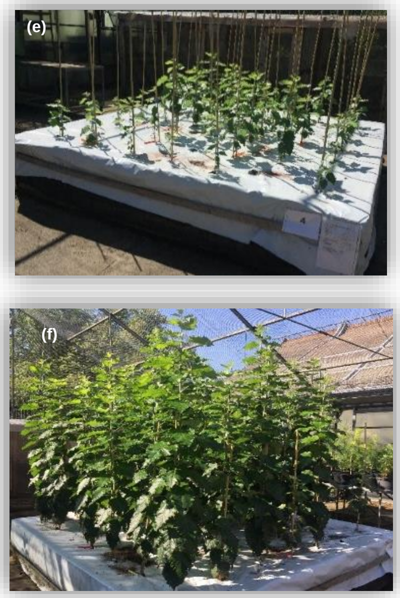


**Supplementary Figure S10. Planting scheme of wildtype and transgenic *Sc*WS expressing poplars (*Populus* × *canescens*) in mixtures under outdoor conditions** (a, b). The poplars were planted in fall 2018. The morphological and physiological measurements started in May 2019 (c,e) and drought treatments started in August 2019 (d,f). Lines used in this experiment, *Sc*WS1 (G1L3), *Sc*WS2 (G1L5.I), *Sc*WS3 (G1L4), *Sc*WS4 (G4L1) and wild type (WT).

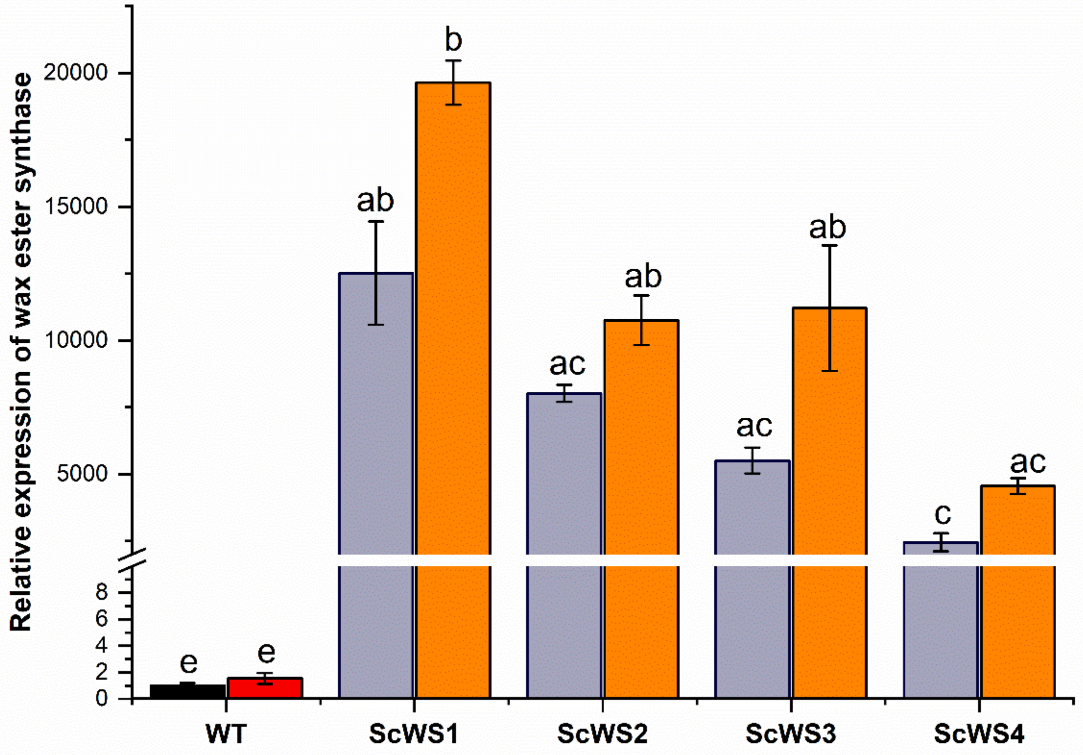


**Supplementary Figure S11. Expression of *Sc*WS and WDS1 (wildtype, WT) in WT and *Sc*WS-expressing *P*. x *canescens* under outdoor conditions**. The expression levels were determined with primers for *Sc*WS and for wax ester synthase/diacylglycerol acyltransferase (WSD1) (cf. Supplement Table S3). Black and grey bars: well-irrigated plants, red and orange bars: drought-stressed plants Leaf number five from the top was harvested on 13th September 2019, 6 weeks after drought exposure and used for the analysis. Data show means (± SE, n = 4 plants per line and treatment). Different letters indicate significant differences among the treatment and lines at P ≤ 0.05 (post hoc Tukey test).

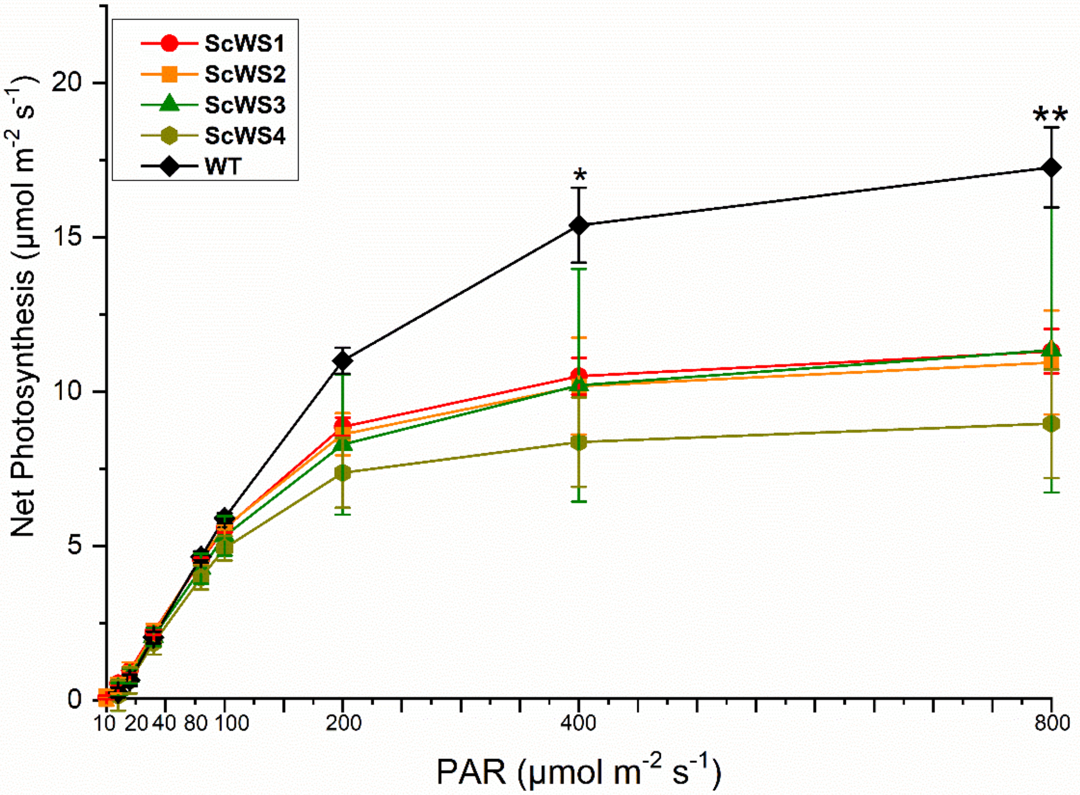


**Supplementary Figure S12. Light response curve of photosynthesis of *Sc*WS lines and wildtype *P.* x *canescens***. Data show means (± SE, n = 3 independent plants per line). PAR, photosynthetically active radiation. Stars denote significant differences: ∗ P ≤ 0.05; ∗∗P ≤ 0.01 of the WT compared with the *Sc*WS lines.

**Supplementary Table S1: Respiration, transpiration and stomatal conductance in darkness of well-irrigated and drought-stressed *Sc*WS lines and wildtype *P*. x *canescens* in a long-term greenhouse experiment**. The measurements were performed at night under the following conditions: air temperature 18.0 ± 0.3 °C, air humidity 57.9 % ± 2.1 %, ambient $CO_2$ 395.6 vpm and photosynthetic photon flux at leaf level 0 µmol $m^{-2}$ $s^{-1}$. Measurements were conducted 12 days after the start of the drought treatment. Data show means (±SE, n = 4). Two-way ANOVA with line and treatment as main factors was used to compare differences of means among poplar lines and drought, followed by a posthoc test (Tukey). Means that differ at p ≤ 0.05 are indicated by different letters.

| Line | Treatment | Respiration (µmol $m^{-2}$ $s^{-1}$) | SE | | Stomata Conductance (mol $m^{-2}$ $s^{-1}$) | SE | | Transpiration (mmol $m^{-2}$ $s^{-1}$) | SE | |
|---|---|---|---|---|---|---|---|---|---|---|
| **ScWS-1** | **Control** | -1.048 | 0.10 | ab | 0.06 | 0.019 | a | 1.73 | 0.53 | a |
| **ScWS-2** | **Control** | -1.04 | 0.15 | ab | 0.038 | 0.012 | ab | 1.178 | 0.33 | ab |
| **WT** | **Control** | -1.28 | 0.27 | a | 0.029 | 0.0083 | a | 0.907 | 0.24 | bc |
| **ScWS-1** | **Drought** | -0.75 | 0.12 | b | 0.017 | 0.004 | b | 0.55 | 0.11 | bc |
| **ScWS-2** | **Drought** | -0.76 | 0.21 | b | 0.016 | 0.0042 | b | 0.53 | 0.12 | bc |
| **WT** | **Drought** | -1.16 | 0.17 | ab | 0.012 | 0.0002 | b | 0.413 | 0.004 | c |

**Supplementary Table S2: Respiration, transpiration and stomatal conductance in darkness of well-irrigated and drought-stressed *Sc*WS lines and wildtype *P.* x *canescens* under field conditions**. The measurements were performed at night under the following conditions: air temperature 18.0 ± 0.3 °C, air humidity 57.9 % ± 2.1 %, ambient $CO_2$ 395.6 vpm and photosynthetic photon flux at leaf level 0 µmol $m^{-2}$ $s^{-1}$. Measurements were conducted 4 weeks after the start of the drought treatment. Data show means (±SE, n = 4). Two-way ANOVA with line and treatment as main factors was used to compare differences of means among poplar lines and drought, followed by a posthoc test (Tukey). Means that differ at p ≤ 0.05 are indicated by different letters.

| Line | Treatment | Respiration (µmol $m^{-2}$ $s^{-1}$) | SE | | Stomata Conductance (mol $m^{-2}$ $s^{-1}$) | SE | | Transpiration (mmol $m^{-2}$ $s^{-1}$) | SE | |
|---|---|---|---|---|---|---|---|---|---|---|
| **ScWS-1** | **Control** | -0.207 | 0.13 | ab | 0.04 | 0.015 | a | 5.93 | 1.29 | ab |
| **ScWS-2** | **Control** | -0.34 | 0.20 | ab | 0.046 | 0.013 | ab | 5.84 | 1.82 | ab |
| **ScWS-3** | **Control** | -0.23 | 0.03 | ab | 0.045 | 0.0097 | a | 5.12 | 0.51 | ab |
| **ScWS-4** | **Control** | -0.28 | 0.09 | ab | 0.04 | 0.009 | ab | 5.86 | 0.31 | ab |
| **WT** | **Control** | -0.44 | 0.25 | a | 0.09 | 0.013 | c | 7.76 | 0.34 | c |
| **ScWS-1** | **Drought** | -0.13 | 0.01 | b | 0.033 | 0.011 | a | 4.14 | 0.69 | ab |
| **ScWS-2** | **Drought** | -0.11 | 0.06 | b | 0.044 | 0.02 | a | 5.76 | 1.13 | ab |
| **ScWS-3** | **Drought** | -0.11 | 0.15 | b | 0.055 | 0.01 | ac | 7.22 | 1.14 | a |
| **ScWS-4** | **Drought** | -0.1 | 0.11 | b | 0.061 | 0.016 | ab | 7..22 | 0.22 | a |
| **WT** | **Drought** | -0.13 | 0.03 | b | 0.087 | 0.02 | bc | 5.55 | 1.45 | b |

**Supplement Table S3: Gas exchange of *P.* x *canescens* wildtype and *Sc*WS lines under outdoor conditions in 2020 (second growth phase).** Gas change measurements were conducted once a week during the drought using LICOR 6800, from 10am to 3pm. n = 5 per line per treatment. The average air temperature and ambiance humidity during the measurement were 23.5 ± 0.3°C and 58% ± 2%. The photon flux density of the photosynthetically active radiation was developed to 800 µmol $m^{-2}$ $s^{-1}$ at the leaf status. WUE was obtained as the ratio of photosynthetic rate to transpiration rate.

| Line | Photosynthesis (µmol $m^{-2}$ $s^{-1}$) | SE | | Stomata conductance (mmol $m^{-2}$ $s^{-1}$) | SE | | Transpiration (mmol $m^{-2}$ $s^{-1}$) | SE | | WUE (µmol $mol^{-1}$) | SE | |
|---|---|---|---|---|---|---|---|---|---|---|---|---|
| **ScWS-1** | 12.01 | 2.47 | ab | 200 | 95 | a | 2.86 | 1.04 | a | 5218.45 | 2651.24 | b |
| **ScWS-2** | 15.51 | 2.93 | ab | 350 | 115 | a | 4.01 | 1.05 | a | 4263.76 | 1717.42 | b |
| **ScWS-3** | 15.11 | 3.57 | ab | 250 | 118 | a | 3.86 | 0.99 | a | 3032.79 | 1150.63 | ab |
| **ScWS-4** | 10.87 | 2.37 | b | 410 | 113 | a | 4.73 | 1.06 | a | 3509.13 | 1173.54 | ab |
| **WT** | 20.79 | 3.69 | ab | 1210 | 200 | b | 6.35 | 1.07 | b | 3585.95 | 1352.22 | a |

**Supplementary Table S4:** List of the primers and Potri numbers for genes used for the cloning (1,2) and for expression analyses by qRT PCR (3). For cloning, primers were designed with 58°C melting temperature using Genious Prime (Biomatters, Ltd., Auckland, New Zealand, https://www.geneious.com). For qRT PCR, primers were designed with the PerlPrimer software version 1.1.20 (https://perlprimer.sourceforge.net, open-source PCR primer design) with maximum of 150 bp and 58-62 °C annealing temperature. *Ptr*PPR_2, and *Ptr*Rpp14 are the housekeeping genes used for the normalization in the q RT PCR experiments.

| | **Gene name** | **Potri Nr.** | **Sequence** |
|---|---|---|---|
| **1** | *Sc*WS | Cloning | FOR: attL1- GGGGACAAGTTTGTACAAAAAAGCAGGCTTAGTCGACATGGAGGTGGAG |
| | | | REV: attL2- GGGGACCACTTTGTACAAGAAAGCTGGGTTCTCACCACCCCAACAAACC |
| **2** | *p*DONR201 | Cloning | FOR: TCGCGTTAACGCTAGCATGGATCTC |
| | | | REV GTAACATCAGAGATTTTGAGACAC |
| **3** | *Sc*WS | | FOR: ATG GTG GTG AAG AAG GCG G |
| | | | REV: TCC AGT CAC CAT CAC GAA CC |
| | CER1 | Potri.014G152300 | FOR: TCTTCATCACCACTATCTCTACTC |
| | | | REV: GAATCACAGAAGTAATGGGCTC |
| | CER2 | Potri.001G319200 | FOR: GTCTTTGTTCAGTTCACTTGGT |
| | | | REV: ATGTGTTGATGAAGGTTGAAGC |
| | CER4 | Potri.004G185000 | FOR: GAATAGTTGATGTGATACCAGCAG |
| | | | REV: CACGGGATTTCTCACAGAGG |
| | CER6 | Potri.008G120300 | FOR: GCACTGGTACTTGTTCAAACTC |
| | | | REV: AATTTGTTCAGATACAGGGAGG |
| | WSD1 | Potri.001G304600 | FOR: TCCATGCAGGATCTATCCGA |
| | | | REV: GATATGATACGCGAAGTGAGCC |
| | WSD4 | Potri.014G098900 | FOR: TTTATCTGTAGTCATCATACGCCC |
| | | | REV: TGCTGGTATCACATCAACTATTCC |
| | OSP1 | Potri.014G160100 | FOR: GTGATGTGCTATCTGTGATACC |
| | | | REV: CCTATGCTTATCATGTGTATACCC |
| | MYB96 | Potri.004G126700 | FOR: AAAGCAAGGGTAGTGTTCTTACTC |
| | | | REV: CTAAGCAAACCTGTATTAGTAGGC |
| | MYB94 | Potri.017G082500 | FOR: CACCTTGCTGTGATAAGATAGGA |
| | | | REV: TCTAAGCAGTCCTGTACTAGTTGG |
| | *Ptr*PPR_2 | Potri.012G141400 | FOR: ATCGTTCCAAGTCAAGTATGTG |
| | | | REV: TCAAGGGAGCAACTTTACAG |
| | *Ptr*Rpp14 | Potri.015G001600 | FOR: GCAATGTGAGGAGTTTAGGG |
| | | | REV: TATTAAATGTCTGTGCTGTAGTGTG |

**Supporting Methods S1: Protocols for poplar transformation, scanning electron microscopy of fresh leaf surfaces and cuticular wax analysis**

### Poplar transformation protocol

The *Sc*WS clone was obtained from Prof. I Feussner (Department for Plant Biochemistry, University of Göttingen, Göttingen, Germany). Primers suitable for the Gateway system™ (Biomatters, Ltd., Auckland, New Zealand, https://www.geneious.com) were used for amplification of the vector (Supplement Table S5). The PCR products were confirmed by electrophoresis, purified using the innuPREP PCRpure Kit (analytik jena AG, Jena, Germany) and cloned into the pDONOR-201 vector (Thermo Fisher Scientific, Waltham, MA, USA) containing the p*35S* promoter to create the gateway pEntry clone. To obtain the binary vector, pEntry::p*35S*::*Sc*WS was cloned into pK7WG using the Gateway system (Invitrogen, Waltham, Massachusetts, USA) and used to transform *Agrobacterium tumefaciens* strain GV3101 pMP90 (BacDive, Braunschweig, Germany). For this purpose, aliquots of competent Agrobacteria (200 µl) were thawed in an ice bath. Then, 1 µl, which contained approximately 100 to 1000 ng of the binary vector DNA, was added to the Agrobacteria and incubated on ice for 5 minutes with occasional stirring. Subsequently, the suspension was frozen in liquid nitrogen for 3 minutes, thawed in a water bath at 37°C and further incubated at 37°C for 5 minutes. Then, 800 µl of YEB (yeast extract broth) medium (without antibiotics) was added the suspension was incubated at 28°C on a shaker (Eppendorf, Köln, Germany) for 2 hours. The suspension was then centrifuged for 2 minutes at 5000 rpm at room temperature. Subsequently, the cells were re-suspended in YEB and plated onto two Petri dishes, containing YEB agar supplemented with the appropriate antibiotic as required.

Agrobacteria transformed with the binary vector were grown over night in 4 ml YEB – medium (with following ingredient: beef extract 5g/l, yeast extract 1g/l, peptone 5g/l, sucrose 5g/l, $MgSO_4$ 0.3 g/l, and agar 20 g/l, pH 7.2) containing antibiotics (Rifampicin, 50 µg $ml^{-1}$; Gentamycin, 25 µg $ml^{-1}$) at 28°C in darkness on an orbital shaking incubator (Sanyo Gallenkamp PLC, Cambridge, United Kingdom) at 90 rpm to generate the starter culture. The starter culture (2 ml) was used to inoculate 100 ml YEB medium (without antibiotics) at 28°C. The culture was grown in darkness at 28°C on a shaker at 90 rpm for about 3 to 5 hours to reach an $OD_{600}$ of 0.3 to 0.5. Then, 20 µM 3´,5´-dimethoxy-4´-hydroxy acetophenone (Sigma Aldrich, Darmstadt, Germany) was added and the incubation was continued for another 30 min.

To transform *P*. x *canescens*, 4-week-old plantlets from a tissue culture were used. The leaves of the plantlets were removed. The stem was divided into small segments of 1 to 2 cm length and transferred into the suspension with the transformed Agrobacteria. The stem segments were incubated in darkness at 28°C for 30 min on a shaker (120 rpm). The stem sections were strained, drained, and transferred to agar plates containing ½ MS (Murashige and Skoog, 1962) medium supplemented with 7g $L^{-1}$ plant agar, 20 g $L^{-1}$

sucrose and were kept in darkness at 25°C for one week. The stem sections, overgrown with Agrobacteria, were washed three times in sterile distilled water with 400 µg $ml^{-1}$ ticarcillin clavulanate (Duchefa, Haarlem, Netherland) for three minutes in an Erlenmeyer flask. Sterile tap water without ticarcillin clavulanate was used for the last washing step.

To screen for the positively transformed plantlets, the stem sections were transferred into selection medium (½MS medium supplemented with 20 g $l^{-1}$ sucrose, and 7 g $l^{-1}$ plant agar, pH 5.8 and the following antibiotica: Pluronic F-68 (0.01%), Thidiazuron (0.0022 mg $l^{-1}$), Cefotaxim (150 mg $l^{-1}$), Timentin (150 mg $l^{-1}$), Kanamycin (50 mg $l^{-1}$) (Duchefa, Haarlem, Netherland). The stem sections were kept for four to six weeks under long day conditions with 16 h of low light, approximately 10 µmol photons $m^{-2}$ $s^{-1}$ photosynthetic active radiation (PAR) (fluorescent lamps L18W/840, Osram, Munich, Germany) at 22°C and 20 to 40 % humidity. When shoots emerged, the plant culture was transferred to rooting medium (½ MS medium, 20 g $l^{-1}$ sucrose, and 7 g $l^{-1}$ plant agar, pH 5.8 with the selection antibiotic Kanamycin (50 mg $l^{-1}$) and incubated under the same conditions above but with 20 µmol photons $m^{-2}$ $s^{-1}$ PAR. Roots produced within five to eight weeks indicated successful transformation. The rooted shoots were grouped in lines and transferred to glass jars (350 ml) with rooting medium with Kanamycin (50 mg $l^{-1}$). Regenerating plantlets from independent transformation events were tested for the presence of the target gene *Sc*WS by Sanger sequencing (Microsynth Seqlab, Göttingen, Germany) after extracting the genomic DNA of one leaf per line (Edwards *et al.*, 1991). Successfully transformed independent lines carrying the *p*35S::*Sc*WS constructs were denominated ScWS1, ScWS2… ScWSn. The lines were kept separately in tissue culture as described by Müller et al. (2013) and were used as the stock for further experiments.

### Scanning electron microscopy of fresh leaves

Two rectangular pieces of approximately 1 cm x 1 cm were cut from each leaf with a double edge razor blade (Wilkinson Sword, United Kingdom) and used for inspection of the ad- and abaxial surface as described by Dreischhoff *et al.*, (2023). The specimen was transferred on top of a Standard SEM Pin Stub Mount (diameter 12.7 mm, Plano GmbH, Wetzlar), which had been covered by conductive double-sided adhesive carbon tape (Plano GmbH, Wetzlar, Germany). Immediately afterwards, the sample was covered with a gold layer of 10 nm thickness (Q150R S/E/ES plus sputter coater, Quorum Technologies Ltd, Lewes, United Kingdom). Preliminary tests showed that preceding freeze-drying can be skipped, if the samples were measured within a short period of time (about 2 days). After sputter coating, the edges of the sample were sealed with a conductive carbon cement (Plano GmbH, Wetzlar, Germany). SEM imagine acquisition was performed in the back-scattered electron mode at a voltage of 15kV and a resolution of 1024 pixels with Phenom ProX (G5) desktop SEM (Phenom-World, Eindhoven, Netherlands). Images of two different leaf area were recorded at magnifications of 350x, 1000x, 2000x, 4000x, and 8000x.

### Cuticular wax analyses

For cuticular wax extraction, three leaf discs of 14 mm in diameter with an area of 2.5 $cm^2$ were collected and analyzed as previously described (Haslam and Kunst, 2013). Plant tissues were immersed in chloroform for 30 s containing 2.5 µg of *n*-tetracosane (C24 alkane) (Merck KGaA, Darmstadt, Germany) as an internal standard. Samples were dried under a stream of nitrogen at room temperature, resuspended in 200 µL of chloroform and evaporated, before derivatization with 10 µL N,O-bis(trimethylsilyl) trifluoroacetamide, 1 % chlorotrimethylsilane (Merck KGaA) and 10 µL of pyridine (Merck KGaA) for 1 h at 80 °C. Derivatization reagents were evaporated under nitrogen gas and samples were resuspended in 40 µL chloroform. For quantitative analysis, 2 µL of each sample were injected onto an Agilent 6890 gas chromatograph coupled with flame ionization detection equipped with a DB1-ht column (30 m x 0.32 mm, 0.1 µm coating thickness [Agilent Technologies Deutschland GmbH, Ratingen, Germany]) using a helium carrier gas inlet at a flow rate of 1.2 mL/min. Cuticular wax analysis was performed in a 5:1 split mode. The oven temperature gradient was as follows: 50 °C for 2 min, ramped by 40 °C/min to 200 °C, held at 200 °C for 1 min, increased by 3 °C/min to 320 °C and held at 320 °C for 15 min. The amount of wax was determined by comparing the area of the internal standard using the Agilent ChemStation software and expressed per unit tissue area ($cm^2$).

For peak identification, the identity of the wax components was confirmed via GC coupled with a mass selective detector (GC-MS) equipped with a DB1-ht column (Agilent 5977B mass selective detector connected to an Agilent 7890B GC system). The mass range was set between 50 and 900 amu. The transfer line was set at 280 °C, the ion source at 230 °C, and the electron energy was 70 eV.
